# SUMO protease and proteasome recruitment at the nuclear periphery differently affect replication dynamics at arrested forks

**DOI:** 10.1101/2023.11.13.566856

**Authors:** Kamila Schirmeisen, Karel Naiman, Karine Fréon, Laetitia Besse, Shrena Chakraborty, Antony M. Carr, Karol Kramarz, Sarah AE Lambert

## Abstract

Nuclear pores complexes (NPCs) are genome organizers, defining a particular nuclear compartment enriched for SUMO protease and proteasome activities, and acting as docking sites for DNA repair. In fission yeast, the anchorage of perturbed replication forks to NPCs is an integral part of the recombination-dependent replication restart mechanism (RDR) that resumes DNA synthesis at terminally dysfunctional forks. By mapping DNA polymerase usage, we report that SUMO protease Ulp1-associated NPCs ensure efficient initiation of restarted DNA synthesis, whereas proteasome-associated NPCs sustain the progression of restarted DNA polymerase. In contrast to Ulp1-dependent events, this last function occurs independently of SUMO chains formation. By analyzing the role of the nuclear basket, the nucleoplasmic extension of the NPC, we reveal that the activities of Ulp1 and the proteasome cannot compensate for each other and affect RDR dynamics in distinct ways. Our work probes the mechanisms by which the NPC environment ensures optimal RDR.

**Highlights:**

● Ulp1-associated NPCs ensure efficient initiation of restarted DNA synthesis, in a SUMO chain-dependent manner
● Proteasome-associated NPCs foster the progression of restarted DNA synthesis, in a SUMO chain-independent manner
● The nucleoporin Nup60 promotes the spatial sequestration of Ulp1 at the nuclear periphery
● Ulp1 and proteasome activities are differently required for optimal recombination-mediated fork restart.

## Introduction

The eukaryotic genome is folded in 3D within a membrane-less compartmentalized nucleus. Nuclear organization constitutes a critical layer of regulation of DNA-associated transactions and an important determinant of genome integrity^1^. The stability of the genome is jeopardized during DNA replication; the progression of the replisome being recurrently threatened by a broad spectrum of obstacles that cause replication fork slowing, temporary fork stalling or terminal fork collapse^2^. Such alterations of fork progression are a hallmark of replication stress. Failure to safeguard genome stability upon replication stress is a potent driving force behind the onset and progression of human diseases including cancer^3^. While multiple replication fork-repair pathways can be engaged at stressed forks to promote the completion of genome duplication, they result in variable outcomes for genome stability and thus must be carefully controlled and regulated. Our current knowledge of the regulatory functions played by nuclear organization in the usage of fork repair pathways remains in its infancy.

Among the fork-repair pathways, homologous recombination (HR) is particularly active in protecting, repairing and restarting stressed forks, making HR an efficient tumor suppressor mechanism^4^. The central factor of the HR machinery is the Rad51 recombinase that forms a nucleoprotein filament on single-stranded DNA (ssDNA), with the assistance of a loader, known as Rad52 in yeast models. In a non-recombinogenic mode, the Rad51 filament limits the degradation of ssDNA by various nucleases, thus ensuring the protection and integrity of stressed forks. In a recombinogenic mode, HR repairs broken forks with a single-ended double-strand break (DSB) by a mechanism called break-induced replication (BIR) and promotes replication resumption at DSB-free collapsed forks by a mechanism called recombination-dependent replication (RDR)^5^. Both BIR and RDR are associated with non-canonical DNA synthesis, approximatively 100 times more mutagenic than canonical replication. Furthermore, during BIR and RDR, both DNA strands are synthesized by DNA polymerase delta (Pol δ)^6,7^. These features allow experimental differentiation between DNA replicated by a repaired/restarted fork and DNA replicated by a canonical origin-born fork. Although stressed forks have the potential to relocate to the nuclear periphery (NP), little is known about the contribution of such changes in nuclear positioning in regulating the replicative functions of the HR machinery.

3D genome folding within the complex nuclear environment is a critical layer of DNA repair regulation. A striking example is the DNA damage response-dependent fate of DSBs that relocate to the NP or shift away from heterochromatin compartments to achieve error-free repair^8,9^. This led to the concept that the membrane-less nuclear compartment exhibits distinct DNA repair capacities and that DNA repair machineries are spatially segregated. Nuclear pore complexes (NPCs) are macromolecular structures embedded in the nuclear envelope (NE) that act as nuclear scaffolds to regulate cellular processes via a wide range of mechanisms^10^. The overall structure of NPCs is conserved among eukaryote kingdom, being composed of multiple copies of 30 different nucleoporins that associate in stable sub-complexes. The core NPC defines a central channel composed of transmembrane and channel nucleoporins. This core complex assembles with the outer and inner rings at the cytoplasmic and nuclear sides, respectively. A Y-shaped structure, located both at the cytoplasmic and nuclear side of NPCs, called in fission yeast Nup107-Nup160 complex, is crucial for NPCs organization and proper segregation of chromosomes in eukaryotes^11–13^. The final composition of individual NPCs is variable, depending on their position within the NE, suggesting that the NPC structure is dynamic. In particular, the nuclear basket, a nucleoplasmic extension of the core NPC, is the most dynamic part and NPCs localized in the nucleolar part of the NE are more frequently devoid of a nuclear basket^12^. The primary function of NPCs is the transport of macromolecules from the cytoplasm to the nucleus and mRNA export. NPCs also define a particular nuclear compartment enriched for the SUMO SENP protease and the proteasome and act as docking sites for DSBs and perturbed replication forks^8^.

Stressed forks can relocate to the NP and, in some cases, anchor to NPCs^14^. These include forks stalled by structure-forming DNA sequences, telomeric repeats, DNA-bound proteins and replication inhibitors^15–21^. Although distinct scenarios arise depending on the source of replication stress and the model organism, the common emerging concept is that nuclear positioning of replication stress sites influences the usage of fork repair pathways. For example, in *Saccharomyces cerevisiae* (Sc), forks stalled within telomeric repeats associate with NPCs to restrict error-prone HR events and maintain telomere length^18^. Forks stalled by CAG repeats, prone to form secondary DNA structures, also anchor to NPCs, in a SUMO-dependent manner^16^. In this instance, SUMOylated RPA on ssDNA at the stalled fork inhibits Rad51 loading, which is permitted only after NPC anchorage that subsequently favors error-free fork restart^17^. Changes in nuclear positioning are far from being a yeast-specific phenomenon. Upon DNA polymerase inhibition, stalled forks in human cells relocate to the NP to minimize chromosomal instability and ensure timely fork restart^20^. Additionally, stressed forks at human telomeres relocate to NPCs to maintain telomere integrity^19^.

We previously reported that, in the yeast *Schizosaccharomyces pombe* (Sp), dysfunctional forks relocate in a SUMO-dependent manner and anchor to NPCs for the time necessary to achieve RDR^15^. This change in nuclear positioning is critical to spatially segregate the subsequent steps of RDR, with dysfunctional forks being processed and remodeled in the nucleoplasm to load Rad51. SUMO chains that are generated by the E3 SUMO ligase, Pli1, trigger relocation to NPCs but limit also the efficiency of HR-mediated DNA synthesis for fork restart. Relocation of dysfunctional forks to NPCs allows SUMO conjugates to be cleared by the SUMO deconjugating enzyme, Ulp1, which is sequestrated at the NP^22^. Therefore, NPCs are an integral part of RDR regulation to promote HR-dependent DNA synthesis at dysfunctional forks. However, the dynamics underlying this process remain unexplored. In particular, the contribution of NPCs to non-canonical Pol δ/Pol δ DNA synthesis, a hallmark of HR-restarted forks, has not been addressed. Here, by mapping DNA polymerase usage during HR-mediated fork restart, we reveal that the SUMO protease, Ulp1, and the proteasome differentially affect the dynamics of HR-dependent fork restart by ensuring efficient DNA synthesis resumption and by sustaining the dynamic progression of the restarted fork, respectively. Moreover, by studying the role of the nuclear basket in RDR, we show that Ulp1 and the proteasome do not compensate for each other, with Ulp1 being critical to counteract the inhibitory effect of SUMO chains but not the proteasome. Our study uncovers mechanisms by which the NPC compartment acts as a critical environment for optimal HR-dependent fork restart.

## Results

To investigate the contribution of the NP to the dynamics of HR-mediated fork restart, we exploited the *RTS1* replication fork barrier (RFB) that promotes the polar arrest of a single replisome at a specific genomic location (Fig. 1a)^23^. The activity of the RFB is fully dependent on the Rtf1 protein that binds to the *RTS1* sequence. The expression of Rtf1 can be artificially regulated by the *nmt41* promoter to allow Rtf1 repression in thiamine-containing media (RFB OFF condition) and its expression upon thiamine removal (RFB ON condition). Alternatively, the *rtf1* gene can be deleted and the results compared with an *rtf1*+ strain. Forks arrested at the RFB become fully dysfunctional and undergo controlled degradation of nascent strands by the end-resection machinery to generate ssDNA gaps of ∼ 1 Kb in length^24,25^. RPA, Rad52 and Rad51 are loaded onto these ssDNA gaps, ensuring fork protection until the arrested fork is either fused with a converging fork or actively restarted by RDR, which occurs approximately 20 minutes after the arrest^6,26–28^. The restarted fork is associated with a non-canonical, mutagenic DNA synthesis in which both strands are synthesized by Pol δ, making it insensitive to the RFB^26,27,29,30^.

**Figure 1:**
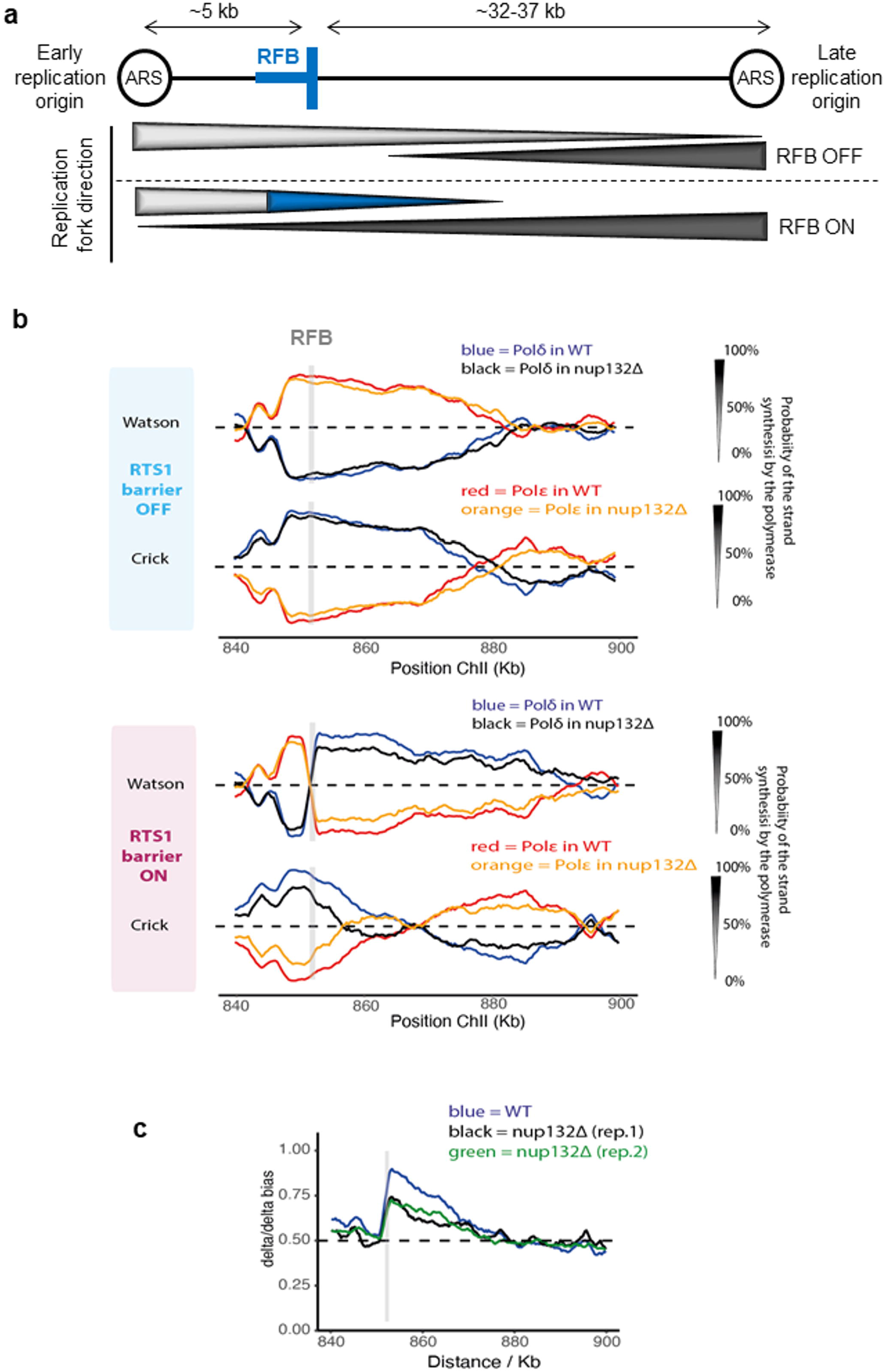
Ulp1-associated NPC promotes the dynamics of recombination-mediated fork restart. **a.** Schematic of the *RTS1*-RFB locus on chromosome II. The position of the *RTS1*-RFB is indicated as thick blue bars. The directional RFB blocks the progression of right-moving forks that initiate from the left autonomously replicating sequence (ARS). The direction of unperturbed (RFB OFF) and perturbed replication (RFB ON) forks is indicated by the thickness of the arrows underneath. Light and dark gray bars indicate the progression of canonical rightward and leftward-moving forks, respectively. The green bar indicates the progression of restarted replication forks mediated by homologous recombination. **b.** Pu-Seq traces of the ChrII locus in *RTS1*-RFB OFF (top panel) and ON (bottom panel) conditions in *WT* and *nup132Δ* strains. The usage of Pol delta (in blue and black for *WT* and *nup132Δ* cells, respectively) are shown on the Watson and Crick strands. The usage of Pol epsilon (in red and orange for *WT* and *nup132Δ* cells, respectively) are shown on the Watson and Crick strands. Note the switch from Pol epsilon to Pol delta on the Watson strand at the RFB site (gray bar), which is indicative of a change in polymerase usage on the leading strand in RFB ON condition. **c.** Graph of Pol delta/delta bias over both strands (Watson and Crick) around the RFB site in *WT* and two independent replicates of *nup132Δ* strains. The gray bar indicates the position of the *RTS*1-RFB.

### Ulp1-associated NPCs ensure the efficient priming of recombination-mediated DNA synthesis

We previously reported that the nucleoporin Nup132, part of the Y complex of NPCs core, promotes RDR in a post-anchoring manner and acts downstream of Rad51 loading^15^. The RDR defect observed in *nup132* null cells is caused by the delocalization of Ulp1 from the NP since its artificial tethering to the RFB restored RDR efficiency. Thus, Ulp1-associated NPCs prime HR-dependent DNA synthesis to ensure efficient RDR, but the dynamics of this process are unknown. To address this, we employed the polymerase usage sequencing (Pu-seq) approach that allows the genome-wide mapping of the usage of Pol δ and polymerase epsilon (Pol ε) during DNA replication^31^. Pu-seq makes use of a pair of yeast strains mutated in either Pol δ or Pol ε that incorporate higher levels of ribonucleotides during DNA synthesis. The mapping of ribonucleotides in a strand-specific manner in strains mutated either for Pol δ or Pol ε allows the genome-wide tracking of polymerase usage. Combined with the *RTS1*-RFB, the Pu-seq method allows monitoring the usage frequency of each polymerase separately on both the Watson and Crick strands when the RFB is either inactive (RFB OFF, in an *rtf1Δ* genetic background) or constitutively active (RFB ON, Rtf1 expressed from the *adh1* promoter to maximize fork arrest efficiency)^26^.

At an inactive barrier site (RFB OFF), replication is canonical and unidirectional coming from an early replication origin (leading strand synthesized by Pol ε and lagging strand synthesized by Pol δ) (Fig 1a-b, top panel). This division of labor between Pol δ and ε changed sharply in an RFB ON strain: at the barrier site, Pol ε in the leading strand is switched to Pol δ during the restart of the blocked fork (Fig 1b, bottom panel). This sharp transition characterizes the efficiency of the restart itself. It means that this creates a bias towards Pol δ on both strands (Watson and Crick) downstream of the *RTS1*-RFB site due to the restart. The Pol δ/d bias describes the time needed for the restart as well as the progression of the restarted fork relative to the canonical convergent fork from a late replication origin^26^. Based on the Pol δ/δ bias (Fig. 1c), we estimated that, when compared to *WT* (*nup132*+) cells, only 60% of the expected number of forks were arrested and restarted in *nup132Δ* cells, while the remaining 40% were either not arrested or were arrested and did not restart before being rescued by an incoming leftward moving canonical fork. The increase in Pol ε usage on the Crick strand for ∼10 Kb downstream of the *RTS1* barrier is indicative of this latter scenario (Fig. 1b). Remarkably, this fork-restart defect is consistent with our previous estimation using a proxy-restart assay that exploits the mutagenic DNA synthesis to provide a genetic readout of RDR efficiency. Using this proxy assay, we reported a nearly two-fold reduction in RDR efficiency in *nup132Δ* cells compared to *WT*^15^. Finally, the relative slope of the Pol δ/δ bias disappearance over distance was similar between the two replicates from *nup132Δ* cells and the *WT* strain, indicating that the forks that succeeded to restart progress with similar speed (Fig. 1c).

### The nuclear basket promotes RDR in a pre- and post-anchoring manner

We next investigated the role of the nuclear basket in dealing with replication stress. The *S. pombe* nuclear basket is composed of 4 non-essential nucleoporins: Nup60 (ScNup60), Nup61 (ScNup2, HsNup50), Nup124 (ScNup1, HsNup153) and Alm1 (ScMlp1/2, HsTPR)^12^. A fifth component is the essential nucleoporin Nup211, a second orthologue of ScMlp1/2 and HsTPR. Some of these components are known to contribute to resistance to DNA damage^32,33^. We confirmed that *alm1Δ* cells were highly sensitive to a wide range of replication-blocking agents and bleomycin-induced DSBs, whereas *nup60Δ* and *nup61Δ* cells exhibited mild sensitivity only to hydroxyurea (HU), a replication inhibitor that depletes dNTP pool (Supplementary Fig. 1a).

To establish if this HU sensitivity correlates with a defect in resuming replication following HU treatment, we arrested cells for 4 hours in 20mM HU and then followed DNA content by flow cytometry upon release into HU-free media. Among nuclear basket mutants, only *nup61Δ* cells displayed a defect in the recovery from HU-stalled forks, a defect similar to the one previously reported for *nup132Δ* cells^15^ (Supplementary Fig. 1b): the *WT* strain reached a G2 DNA content 45 minutes after release, whereas both *nup132Δ* and *nup61Δ* cells exhibited an additional 15 minutes delay. This observation is supported by the analysis of chromosomes by Pulse Field Gel Electrophoresis (PFGE). HU treatment prevented chromosomes from migrating into the gel because of replication intermediates accumulation (Supplementary Fig. 1c). *WT* chromosomes migrated into the gel, with twice intensity of an asynchronous culture, 90 minutes after release, indicating complete genome duplication *WT* genome and recovery from HU-stalled forks (Supplementary Fig. 1d). Consistent with the flow cytometry data, only chromosomes from *nup132Δ* and *nup61Δ* cells showed a clear delay in their ability to migrate into the gel and to fully duplicate, confirming a role for Nup61 in promoting DNA replication upon transient fork stalling by HU.

To establish the role of the nuclear basket in promoting replication resumption at the RFB, we first measured replication slippage (RS) downstream of *RTS1*, the proxy measure of non-canonical replication resulting from RDR^30^ (Fig. 2a). The absence of Nup60 and Alm1, but not Nup124 or Nup61, led to a ∼2-fold reduction in the frequency of RFB-induced RS, indicating a reduced RDR efficiency (Fig. 2b). Analysis of replication intermediates by bi-dimensional gel electrophoresis (2DGE) showed that fork arrest and the formation of large ssDNA gaps (>100 bp) at the RFB (which are visualized as a specific “tail” DNA structure emanating from the fork arrest signal and descending toward the linear arc; see red arrow on Fig. 2c)^28^ were unaltered in all four non-essential nucleoporin mutants (Fig. 2c-d). This indicates that the controlled degradation of nascent strands and Rad51-dependent fork protection are unaffected. Thus, the RDR defect observed in *nup60Δ* and *alm1Δ* is not related to defects in the early steps of RDR, from ssDNA gap formation to Rad51 loading.

**Figure 2:**
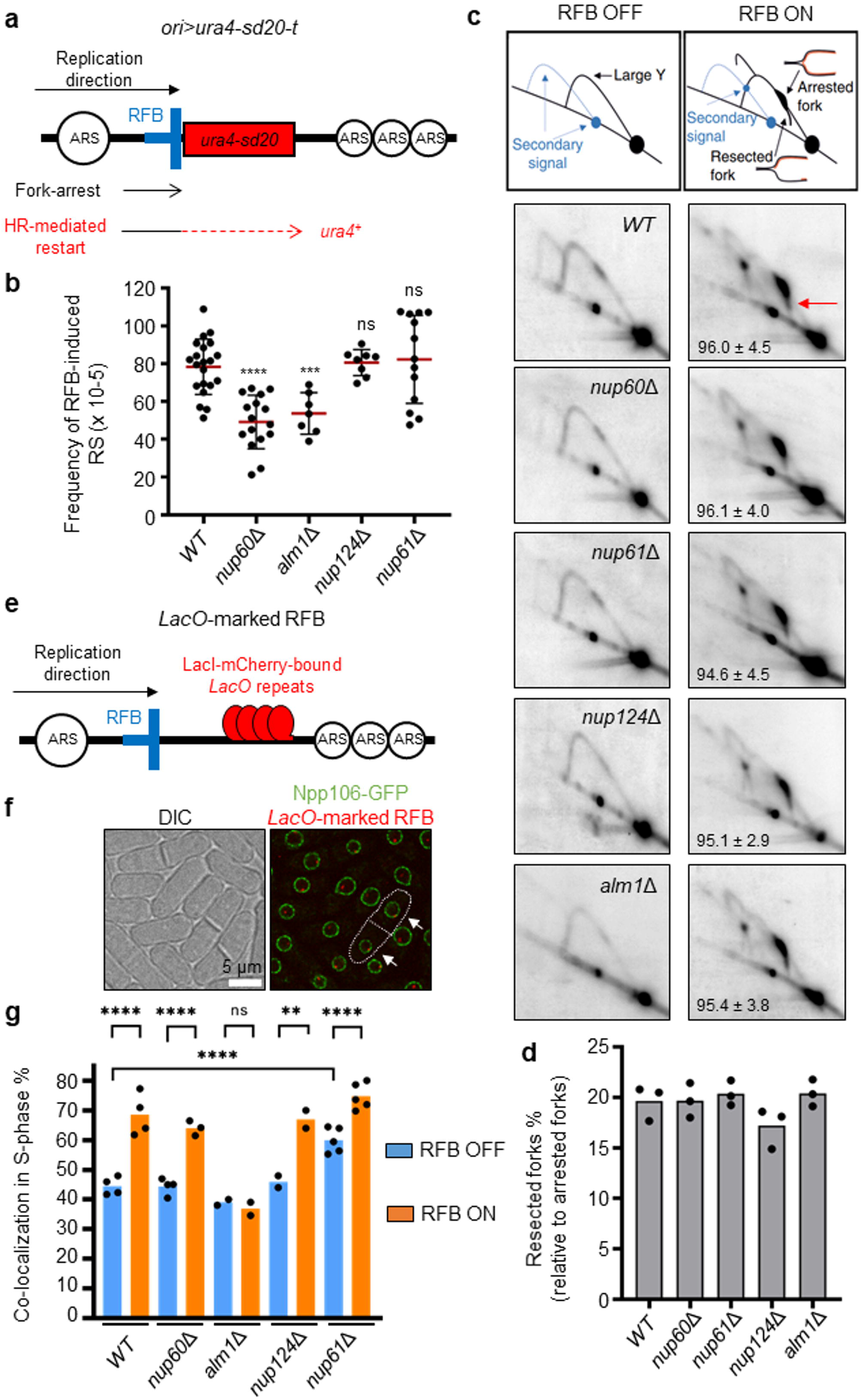
The nuclear basket promotes recombination-dependent replication in a pre- and post-anchoring manner. **a.** Diagram of the *ori>ura4-sd20-t* construct on chromosome III (ori: replication origin, >: *RTS1*-RFB orientation that block right-moving forks, t: telomere). The non-functional *ura4-sd20* allele (red box), containing a 20-nt duplication flanked by micro-homology, is located downstream of the RFB (blue bar). During HR-mediated fork restart, the *ura4-sd20* allele is replicated by an HR-associated DNA synthesis that is liable to replication slippage (RS) resulting in the deletion of the duplication and the restoration of a functional *ura4^+^*gene^30^. ARS: autonomously replicating sequence. **b.** Frequency of RFB-induced RS in indicated strains. Each dot represents one sample from independent biological replicate. Red bars indicate mean values ± standard deviation (SD). *p* value was calculated by two-sided *t*-test (**** *p* ≤0.0001; *** p≤0.001; ns: non-significant). **c.** Top panel: scheme of replication intermediates (RI) analyzed by neutral-neutral 2DGE of the *AseI* restriction fragment in RFB OFF and ON conditions. Partial restriction digestion caused by psoralen-crosslinks results in a secondary arc indicated on scheme by blue dashed lines. Bottom panels: representative RI analysis in indicated strains and conditions. The *ura4* gene was used as a probe. Numbers indicate the % of forks blocked by the RFB ± standard deviation (SD). The red arrow indicates the tail signal resulting from resected forks. **d.** Quantification of resected forks in indicated strains. Dots represent values obtained from independent biological experiments. No statistical difference was detected between the samples using the two-sided *t*-test. **e.** Diagram of the *LacO*-marked RFB. *LacO* arrays bound by mCherry-LacI (red ellipses) are integrated ∼7 kb away from the *RTS1*-RFB (blue bar). **f.** Example of fluorescence (right panel) and bright-field images (left panel, DIC) cells expressing the endogenous Npp106-GFP fusion protein and harboring the *LacO*-marked RFB. Mono-nucleated cells and septated bi-nucleated cells correspond to G2 and S-phase cells, respectively. White arrows indicate co-localization events in S-phase cells. Scale bare: 5µm. **g.** Quantification of co-localization events, shown in f, in S-phase cells in indicated conditions and strains. Dots represent values obtained from independent biological experiments. At least 100 nuclei were analyzed for each strain and condition. Fisher’s exact test was used for group comparison to determine the *p* value (****p≤0.0001; ** p≤0.01; ns: non-significant).

We next investigated the ability of the RFB to relocate to the NP. We employed a strain harboring a *LacO*-marked RFB, expressing LacI-mCherry (Fig. 2e) and an endogenously GFP-tagged Npp106, a NPCs component, to mark the NP (Fig. 2f). We counted co-localization events between the NP and the *LacO*-marked RFB, visualized by a LacI-mCherry focus (see white arrows on Fig. 2f), as previously reported^15^. When the RFB was inactive (RFB OFF), LacI-foci co-localized with the NP in ∼45 % of S-phase cells (Fig. 2g). Upon activation of the RFB (RFB ON), the *LacO*-marked RFB was more often (∼70 %) localized at the NP in *WT* cells^15^. This shift of the active RFB to the NP was observed in all nuclear basket mutants except *alm1Δ* (Fig. 2g). The *nup61Δ* cells exhibited a slight increase in the frequency of co-localization in RFB OFF condition but reached a similar enrichment at the NP to *WT* cells in RFB ON condition. Thus, Alm1 and Nup60 promote RDR in a pre- and post-anchoring manner, respectively.

### The nuclear basket promotes the sequestration of Ulp1 at the nuclear periphery

In budding yeast, several components of the nuclear basket are critical for peripheral Ulp1 localization, including ScNup60 and the synergistic action of ScMlp1 and ScMlp2^34,35^. We thus investigated the expression and the nuclear sub-localization of Ulp1 upon loss of nuclear basket functionality. Ulp1 was C-terminally tagged with GFP and its functionality was established using resistance to genotoxins (Supplementary Fig. 2a). We observed that, in *nup60Δ* and *nup132Δ* mutants, Ulp1-GFP levels were largely abrogated whereas a ∼75 % and ∼60 % reduction was observed in *nup124Δ* and *alm1Δ* backgrounds, respectively (Fig. 3a). Treating cells with bortezomib, a proteasome inhibitor^36^, partly restored Ulp1-GFP protein level in *nup132Δ* and *nup60Δ* cells, similarly to previous findings in budding yeast^34^. However, the sequestration of Ulp1-GFP at the NP was not restored (Supplementary Fig. 2b-c).

**Figure 3:**
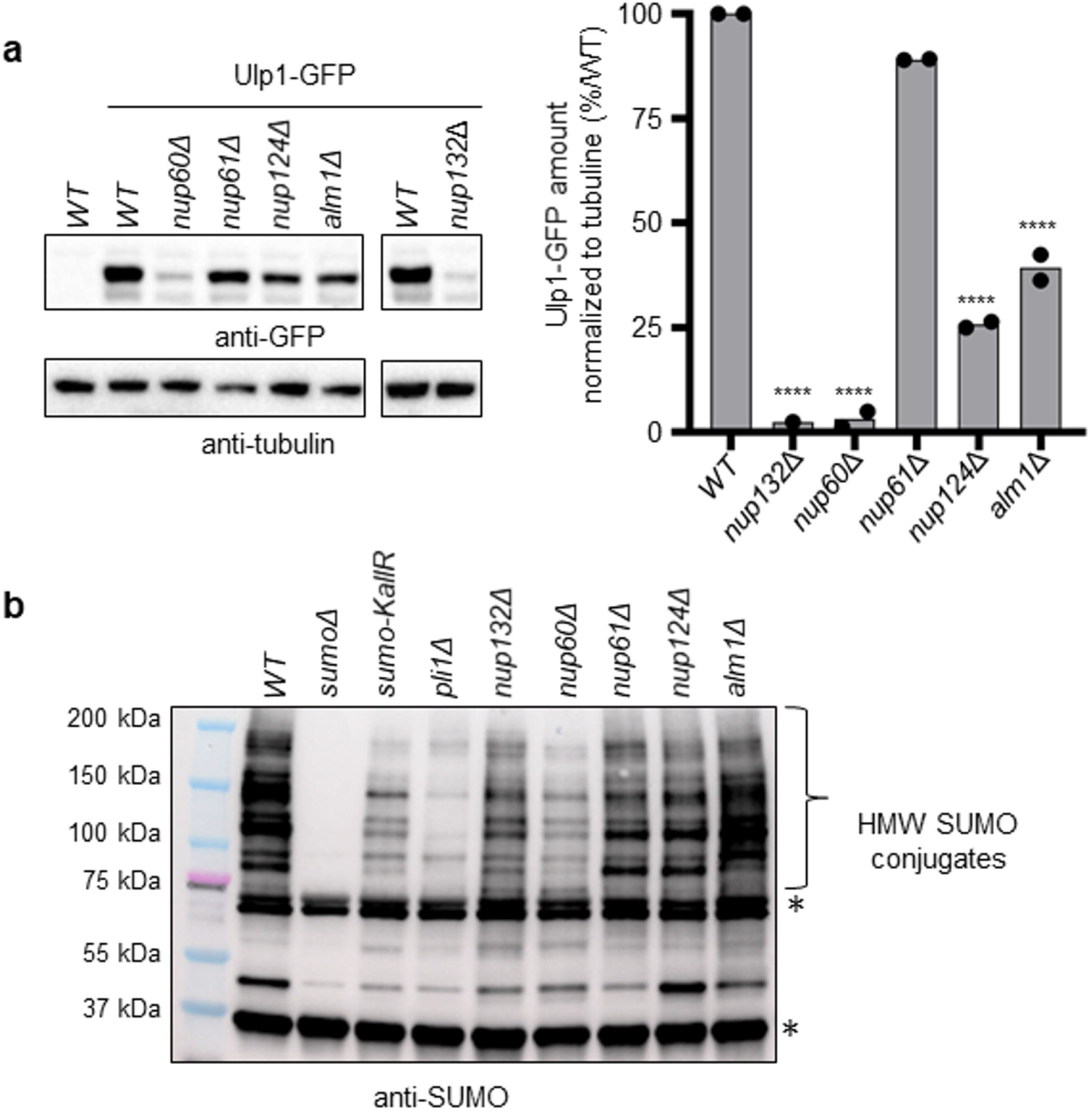
The nuclear basket regulates the expression of the SUMO SENP protease Ulp1. **a.** Left panel: expression of Ulp1-GFP in indicated strains. An untagged *WT* strain was included as control for antibody specificity. Tubulin was used as a loading control. Right panel: quantification. The normalized amount of Ulp1 was calculated by dividing the GFP signal by tubulin signal. The normalized amount of Ulp1-GFP in the mutants is indicated as a percentage of the *WT*. Dots represent values obtained from independent biological experiments. *p* value was calculated by two-sided *t-*test (**** p≤0.0001). **b.** Expression of SUMO conjugates in indicated strains. A strain deleted for *pmt3* gene that encodes the SUMO particle (*sumoΔ*) was added as control for antibody specificity. * indicates unspecific signal.

In *S. pombe*, Ulp1 delocalization leads to the degradation of SUMO chain-modified Pli1, an E3 SUMO ligase, resulting in a global decrease of SUMO conjugates^37^. Consistent with Ulp1 expression being severely lowered and delocalized from the NP in *nup132Δ* and *nup60Δ* (Fig. 3a and Supplementary Fig. 2b), we observed a global reduction in the accumulation of SUMO conjugates, compared to *WT* (Fig. 3b). The pattern of SUMO conjugates in *nup132Δ* and *nup60Δ* backgrounds was similar to the one observed in a strain expressing SUMO-KallR, in which all internal lysines are mutated to arginines to prevent SUMO chains formation^15^. The accumulation of SUMO conjugates was more adversely affected in the absence of Pli1 than in *nup132Δ* and *nup60Δ* cells, suggesting that Pli1 conserves some activity in these genetic backgrounds, as reported for *nup132Δ* cells^15^. Despite a reduced Ulp1 expression in *nup124Δ* and *alm1Δ* cells, the pattern of SUMO conjugates was less affected, suggesting that the remaining Ulp1 sequesters properly at the NP in these genetic backgrounds (Fig. 3b).

To better assign the nuclear basket function in sequestrating Ulp1 at the NP, we employed live cell imaging to detect simultaneously Ulp1 in *WT* and mutant backgrounds and quantify Ulp1 density at the NP. To ensure accuracy, we mixed an equal amount of exponentially growing *WT* cells expressing Ulp1-GFP with *WT* or nuclear basket mutants co-expressing Ulp1-GFP and Cut11-mCherry (Fig. 4a). This approach allowed us to distinguish *WT* cells from the mutated strains within the same microscopy field, and thus accurately quantify peripheral Ulp1 irrespective of exposure and acquisition parameters. In addition, as Cut11 is a transmembrane core NPC nucleoporin, we also could quantify the total amount and density of NPCs. As previously reported^38^, the nuclear morphology of *alm1Δ* cells was different from *WT*, with an increase in nuclear perimeter and size (Supplementary Fig. 3a). The total amount of peripheral Ulp1 decreased in *nup132Δ*, *nup60Δ* and *nup124Δ* cells when compared to *WT* (Supplementary Fig. 3b), resulting in a reduced peripheral Ulp1 density (Fig. 4a-b). Although the total amount of peripheral Ulp1 was slightly increased in *alm1Δ* cells (Supplementary Fig. 3b), the increased nuclear size led to a significant reduction in terms of peripheral Ulp1 density (Fig. 4b). The total amount of Cut11 was variable in all strains when compared to *WT* (Supplementary Fig. 3c) but we observed a clear reduction in peripheral Cut11 density in *alm1Δ* cells because of an increased nucleus size (Fig. 4c). Finally, we quantified co-localization between Cut11-mCherry and Ulp1-GFP signal as a read-out of Ulp1-associated NPCs, using Manders overlap coefficient (Fig. 4d-e) and Pearson correlation coefficient (Supplementary Fig. 3d). As a control, we first assigned co-localization between Cut11-mCherry and Npp106-GFP, two core components of NPCs. Between 80 to 90 % of Cut11 signal was associated with Npp106 under our microscopy conditions, validating our methodological approach (Fig. 4d-e and Supplementary Fig. 3d). In the absence of either Nup132 or Nup60, the low level of Ulp1 appeared to not overlap with Cut11 at the resolution achieved on the images, indicating that Ulp1-associated NPCs are abolished. Despite a lower NPCs density and a reduced Ulp1 expression in the absence of Alm1, Ulp1-associated NPCs were only moderately affected (∼70 % compared to ∼75 % in the *WT* background). In contrast, only ∼50 % of Cut11 signal was correlated with Ulp1 in *nup124Δ* cells (Fig. 4e and Supplementary Fig. 3d), indicating that Ulp1-associated NPCs are less abundant. We concluded that Nup60 and, to a lesser extent, Nup124, are two key components of the nuclear basket that sequester Ulp1 at the NP.

**Figure 4:**
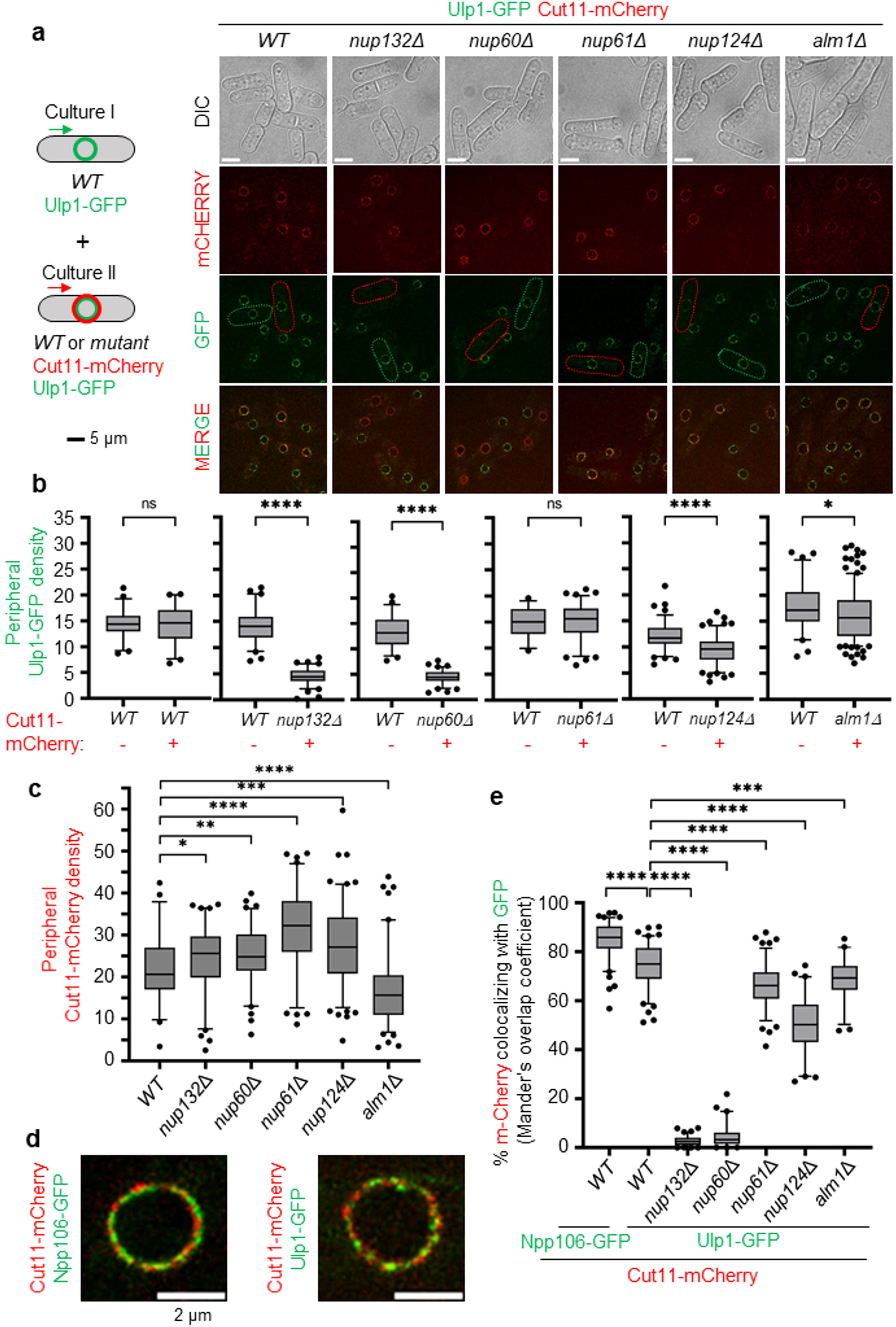
The nuclear basket contributes to sequester the SUMO SENP protease Ulp1 at the nuclear periphery. **a.** Left panel: scheme of the strategy employed by equally mixing two genetically distinct cell cultures. Right panel: representative cell images of Cut11-mCherry and Ulp1-GFP in indicated strains. Green and red cell borders indicate cells from culture I (expressing Ulp1-GFP) and culture II (expressing Ulp1-GFP Cut11-mCherry), respectively. Scale bare 5 µm. **b.** Box-and-whisker plots of Ulp1-GFP density (mean fluorescence intensity) at the nuclear periphery in indicated strains and conditions. Boxes represent the 25/75 percentile, black lines indicate the median, the whiskers indicate the 5/95 percentile and dots correspond to minimum and maximum values. *p* value was calculated by Mann-Whitney U test (**** *p* ≤0.0001; *** p≤0.001; ** p≤0.01; * p≤0.05; ns: non-significant). At least 50 nuclei were analyzed for each strain. **c.** Box-and-whisker plots of Cut11-mCherry density (mean fluorescence intensity) at the nuclear periphery in indicated strains. Boxes represent the 25/75 percentile, black lines indicate the median, the whiskers indicate the 5/95 percentile and dots correspond to minimum and maximum values. *p* value was calculated by Mann-Whitney U test (**** *p* ≤0.0001; *** p≤0.001; ** p≤0.01; * p≤0.05; ns: non-significant). At least 50 nuclei were analyzed for each strain. **d.** Example of the localization of Npp106-GFP and Cut11-mCherry (left panel) or Ulp1-GFP and Cut11-mCherry (right panel) on overlay images. Scale bare: 2 µm. **e.** Box-and-whisker plots of co-localization between Cut11-mCherry and Ulp1-GFP (Mander’s overlap coefficient) in indicated strains. The co-localization between the Npp106-GFP, an inner ring nucleoporin of NPC, and Cut11-mCherry, was performed as a control to show maximum correlation between intensities of those both proteins at the resolution achieved on the images. Boxes represent the 25/75 percentile, black lines indicate the median, the whiskers indicate the 5/95 percentile and dots correspond to minimum and maximum values. *p* value was calculated by Mann-Whitney U test (**** *p* ≤0.0001; *** p≤0.001)

In budding yeast, Mlp1 and Mlp2 act synergistically to sequester Ulp1 to the NP^35^. We therefore addressed the role of the second TPR orthologue Nup211, an essential nucleoporin in *S. pombe*. We employed an auxin-inducible degron (AID) approach using the recently developed AID2 version that makes use of OsTIR1-F74A to target AID-tagged proteins for degradation^39^. Nup211-HA-mAID was efficiently degraded 30 minutes after the addition of 5-adamantyl-IAA and no degradation was observed in the absence of TIR1-F74A (Supplementary Fig. 4a). We observed a ∼40 % reduction in Ulp1-GFP expression 60 minutes after 5-adamantyl-IAA addition, compared to the control strain in which TIR1-F74A is not expressed (compare lines 3 and 4 in Supplementary Fig. 4b). However, Ulp1-GFP expression was slightly decreased in the strain expressing TIR1-F74A in the absence of 5-adamantyl-IAA (compare lines 1 and 2 in Supplementary Fig. 4b). Consistently, these strains showed a significant growth defect when plated on media free of drug (Supplementary Fig. 4c), indicating that either the AID2 system applied to Nup211 is leaky or that the C-terminal degron tag partially compromised Nup211 function. When we quantified peripheral Ulp1-GFP by live-cell imaging, we observed that the addition of 5-adamantyl-IAA led to an increased density of peripheral Ulp1 in *WT* cells and no changes were observed upon degradation of Nup211 (Supplementary Fig. 4d). We concluded that Nup211 makes little contribution to Ulp1 expression and peripheral sequestration. We wanted to test the possibility that Alm1 and Nup211 act synergistically to regulate Ulp1 expression and localization, but we failed in generating viable spores combining *alm1* deletion with the *nup211-HA-mAID* locus.

### Tethering of the RFB to Ulp1-associated NPCs rescues RDR defect in *nup60****Δ*** but not *alm1****Δ*** *cells*

We previously established that SUMO chains trigger relocation of the RFB to NP but also impede HR-mediated DNA synthesis at arrested forks in the absence of Nup132^15^. Thus, when the active RFB shifts to the NP and Ulp1 is no longer recruited at NPCs to degrade SUMO conjugates, RDR is impeded. To test if the same scenario occurs in the absence of Nup60, we employed a previously successful approach to tether Ulp1-LexA to the RFB harboring 8 LexA binding sites (either *t-LacO-ura4:LexBS<ori* for nuclear positioning (Fig. 5a) or *t-ura4-sd20:lexA<ori* for RFB-induced RS^15^ (Fig. 5a). In *WT* cells, the *LacO*-marked RFB was constitutively enriched at the NP upon expression of Ulp1-LexA, whatever its activity (OFF or ON), showing that Ulp1 is successfully tethered to the RFB (Fig. 5a). Consistent with the role of Nup60 in sequestrating Ulp1 at the NP, the inactive RFB did not shift to the NP in *nup60Δ* cells but was efficiently enriched at the NP in RFB ON condition, confirming that Ulp1 is dispensable for anchorage (Fig. 5a). Remarkably, tethering Ulp1-LexA to the active RFB, anchored to NPCs, resulted in an increased frequency of RFB-induced RS in *nup60Δ* cells, indicating that the lack of Ulp1-associated NPCs is a limiting step in promoting HR-mediated DNA synthesis (Fig. 5b). In addition, we combined the *nup60* deletion with SUMO-KallR, which allows only mono-SUMOylation to occur (c.f. Fig. 3b). As previously reported^15^, we observed a slight increase in RFB-induced RS in SUMO-KallR strain, indicating that SUMO chains limit RDR efficiency (Fig. 5c). As expected, preventing SUMO chains in *nup60Δ* cells restored RFB-induced RS to *WT* level, further confirming that Ulp1-associated NPCs are required to overcome the inhibitory effect of SUMO chains on HR-mediated DNA synthesis (Fig 5c).

**Figure 5:**
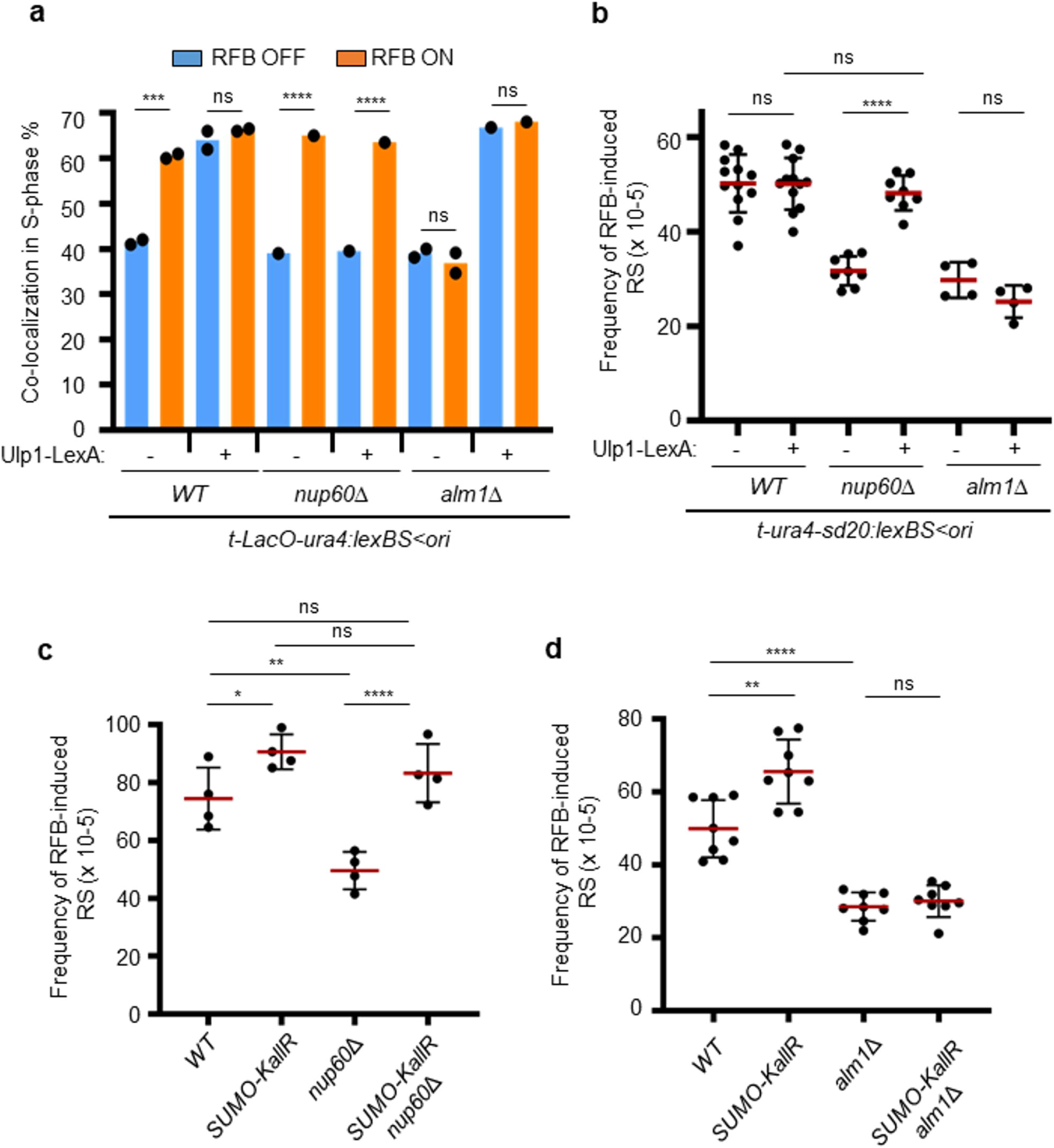
Tethering of the RFB to Ulp1-associated NPCs rescues RDR defect in *nup60Δ* but not *alm1Δ* cells. **a.** Quantification of co-localization events in S-phase cells in indicated conditions and strains. Dots represent values obtained from independent biological experiments. At least 100 nuclei were analyzed for each strain and condition. *p* value was calculated by two-sided Fisher’s exact test (**** p≤0.0001; *** p≤0.001; ns: non-significant). **b and c.** Frequency of RFB-induced RS in indicated strains and conditions. Dots represent values obtained from independent biological experiments. Red bars indicate mean values ± SD. *p* value was calculated by two-sided *t-*test **** *p* ≤0.0001; ** p≤0.01,* p≤0.05 ns: non-significant).

Surprisingly, applying similar approaches to *alm1Δ* cells resulted in different outcomes, indicating a distinct scenario of RDR defect. Preventing SUMO chains formation did not rescue the RDR defect observed in the absence of Alm1 (compare *alm1Δ* and *alm1Δ* SUMO-KallR on Fig. 5d), indicating that this mutant does not suffer from the toxicity of SUMO chains against HR-mediated DNA synthesis. Moreover, tethering Ulp1 to the RFB did not rescue the RDR defect (Fig. 5b). The analysis of the nuclear positioning of the *LacO*-marked RFB showed that the RFB was efficiently shifted to the NP in *alm1Δ* cells regardless its activity, thus allowing bypassing the role of Alm1 in locating the active RFB at the NP (compare RFB ON condition with or without Ulp1-LexA in *alm1Δ* on Fig. 5a). In other words, the artificial anchorage of the RFB to Ulp1-associated NPCs is not sufficient to rescue the RDR defect of *alm1Δ* cells. This indicates that the lack of RFB relocation to the NP is not the underlying cause of the RDR defect and that Alm1 is probably required at NPCs to promote RDR, independently of SUMO chains. Interestingly, Daga and colleagues have reported that Alm1 is required for proper localization of the proteasome to the NE. Several proteasome subunits and anchors, such as Mts2, Mts4 and Cut8, are not properly localized at the NP in *alm1Δ* cells^38^. Interestingly, we previously proposed that RFB relocation to NPCs allows also access to the proteasome to promote RDR^15^. Given the technical difficulty to restore a stoichiometric proteasome at the NP in *alm1Δ* cells, we turned our attention to a viable proteasome mutant to address its role in the dynamic of RDR.

### Proteasome-associated NPCs sustain the dynamic of HR-restarted fork

We previously reported that, in the absence of the proteasome subunit Rpn10, the active RFB shifts to the NP but RDR efficiency was severely decreased^15^. Rpn10 is located at the NP and is a regulatory subunit of the 19S proteasome that physically interacts with Mts4/Rpn1^40–42^. Rpn10 acts as an ubiquitin receptor of the proteasome and its absence results in the accumulation of ubiquitin conjugates. Despite an accumulation of SUMO conjugates in *rpn10Δ* cells (Fig. 6a), we observed that the defect in RFB-induced RS was not rescued by preventing SUMO chains formation (Fig. 6b), indicating a role of the proteasome in promoting RDR independently of counteracting the inhibitory effect of SUMO chains. To probe this function, we applied the Pu-Seq approach to the *rpn10* mutant to compare DNA polymerase usage at and downstream from the barrier site. Based on the Pol δ/δ bias (Fig. 6c-d), we estimated that when compared to *rpn10*+ cells, approximately 85% of the expected number of forks were arrested and restarted in *rpn10Δ* cells (Fig. 6c). Remarkably, the relative slope of the Pol δ/δ bias disappearance over distance was much steeper in the two replicates from *rpn10Δ* cells than in the *WT* strain, indicating a lower speed or increased instability of restarted forks (Fig. 6d). This slow/unstable replication accounts for the increased number of leftward moving canonical forks evident in the Pu-seq traces (Fig. 6c). We estimated that approximately half of restarted forks progress approximately one third of the distance of *WT* restarted forks. This scenario contrasts with that observed in the *nup132Δ* cells, in which fewer forks were restarted but the progression of those that did restart was unaffected. We concluded that both the proteasome and Ulp1 are required at the NP to foster the dynamics of HR-mediated DNA synthesis by affecting, respectively, the efficient initiation of restarted DNA synthesis and the progression of the restarted fork.

**Figure 6:**
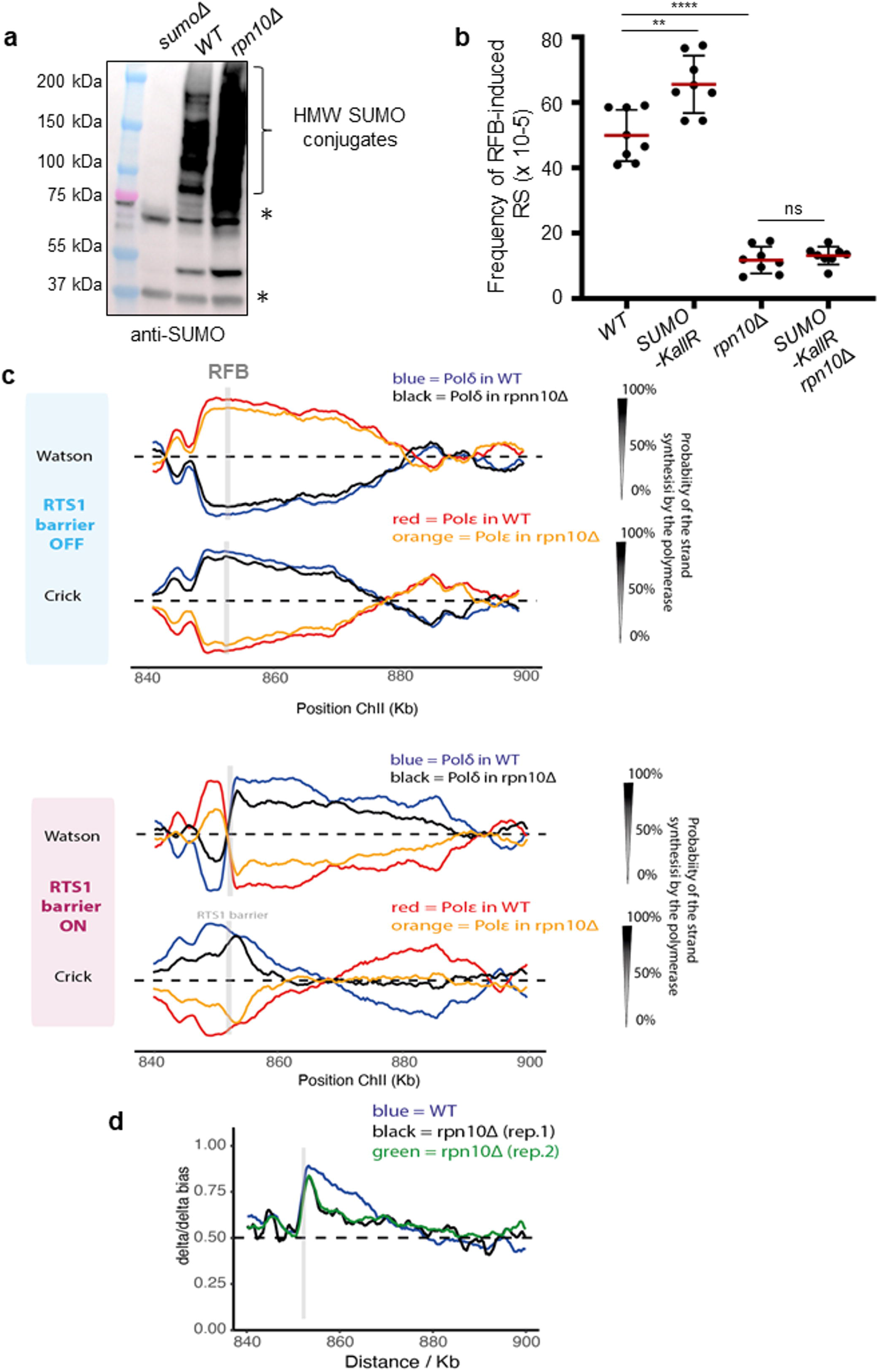
Proteasome-associated NPCs promote the progression of restarted fork. **a.** Expression of SUMO conjugates in indicated strains. A strain deleted for *pmt3* gene that encodes the SUMO polypeptide (*sumoΔ*) was added as control for antibody specificity. * indicates unspecific signal. **b.** Frequency of RFB-induced RS in indicated strains and conditions. Dots represent values obtained from independent biological experiments. Red bars indicate mean values ± SD. *p* value was calculated by two-sided t-test (**** *p* ≤0.0001; ** p≤0.01; ns: non-significant). **c.** Pu-seq traces of the ChrII locus in *RTS1*-RFB OFF (top panel) and ON (bottom panel) conditions in *WT* and *rpn10Δ* strains. The usage of Pol delta (in blue and black for *WT* and *rpn10Δ* cells, respectively) are shown on the Watson and Crick strands. The usage of Pol epsilon (in red and orange for *WT* and *rpn10Δ* cells, respectively) are shown on the Watson and Crick strands. Note that the switch from Pol epsilon to Pol delta on the Watson strand at the RFB site (gray bar) is indicative of a change in polymerase usage on the leading strand in RFB ON condition. **d.** Graph of Pol delta/delta bias in RFB ON condition according to chromosome coordinates in *WT* and two independent replicates of *rpn10Δ* strains. The gray bar indicates the position of the *RTS*1-RFB.

## Discussion

Halted replication forks are diverted to the NP and can associate with NPC components to engage error-free DNA repair pathways^8,14–21^ . How the NPC environment acts mechanistically to foster the dynamics of DNA repair pathways remains unclear. Here, we reveal that NPCs define a particular nuclear compartment that favors the dynamic of HR-dependent DNA synthesis at dysfunctional forks by two distinct mechanisms. The SUMO protease Ulp1 ensures an efficient initiation of restarted DNA synthesis by alleviating the inhibitory effect of SUMO chains. This mechanism requires the sequestration of Ulp1 at the NP which is coordinated by the Y complex and the nuclear basket nucleoporin Nup60. The second mechanism relies on the ability of the nuclear basket to enrich proteasome components at the NP^38,42^, to foster the progress of restarted DNA polymerases. Surprisingly, this last function cannot be bypassed by preventing SUMO chains formation. We establish that Ulp1 and the proteasome affect differently the dynamics of HR-mediated DNA synthesis without compensating for each other.

We previously reported that the Y complex nucleoporin Nup132 promotes RDR in a post-anchoring manner, downstream of Rad51 loading at dysfunctional forks, by sequestrating Ulp1 at the NP to alleviate the inhibitory effect of SUMO chains on HR-mediated DNA synthesis^15^. Here, we reveal that the nuclear basket contributes to this pathway. Akin to budding yeast ^34^, the sequestration of Ulp1 at the NP in *S. pombe* requires the nuclear basket nucleoporin Nup60. Despite our efforts, we cannot rule out a synergistic functions of TPR homologs, Alm1 and Nup211, in the spatial segregation of Ulp1 at the NP. By mapping DNA polymerase usage during HR-dependent fork restart^26^, we establish that Ulp1-associated NPCs are necessary to ensure efficient initiation of restarted DNA synthesis that is likely inhibited by Pli1-dependent formation of SUMO chains of unknown targets. In budding yeast, a similar inhibitory effect of SUMO chains on DNA replication initiation at origins has been reported^43^. The MCM helicase and other replication factors were identified as SUMO chains-modified substrates for targeting by the SUMO protease Ulp2 and proteasomal degradation. Although we did not formally address the function of t SpUlp2 in RDR, our data clearly highlight a role for Ulp1-associated NPCs in counteracting the inhibitory effect of SUMO chains on the initiation of restarted DNA synthesis. Protein-protein docking studies predicted a higher affinity of SpUlp1 towards SUMO particles compared to ScUlp1, suggesting a more specific role of SpUlp1 in targeting SUMO chains than Ulp2^44^. Moreover, the abundance of Ulp1-associated NPCs is not a limiting factor in promoting RDR, as their reduction by 40 % in *nup124Δ* cells does not lead to no noticeable RDR defect.

We previously reported that the proteasome, whose activity is enriched at the NP^42^, promotes RDR in a post-anchoring manner^15^. The mapping of DNA polymerase usage during HR-dependent fork restart reveals that a proteasome defect affects more severely the progression of restarted DNA polymerases, with a reduction of forward movement by up to 70 %, than the initiation of restarted DNA synthesis. This contrasts with Ulp1 function in contributing primarily to the initiation of DNA synthesis with no apparent contribution to the dynamic progression of restarted DNA polymerases. This division of labour between the proteasome and the SUMO protease in ensuring the dynamics of HR-dependent fork restart is reinforced by the fact that these activities cannot compensate for each other. Indeed, the artificial tethering of the RFB to NPCs in the *alm1Δ* mutant shows that Ulp1-associated NPCs are insufficient to promote efficient RDR without a functional proteasome at the NP. Moreover, our genetic analysis establishes that the role of the proteasome in fostering the progress of restarted DNA synthesis is not related to counteracting the inhibitory effect of SUMO chains. This suggests distinct specificities between the proteasome and Ulp1 towards SUMOylated targets which affect differently the dynamics resumption of DNA synthesis at dysfunctional forks. We do not exclude that SUMO-independent poly-ubiquitination, targeted by Rpn10 for proteasomal degradation, plays a role in promoting RDR. However, we previously identified that the SUMO Targeted Ubiquitin Ligase (STUbL) Slx8-Rfp1-Rfp2, a family of E3 ubiquitin ligases that targets SUMOylated proteins for degradation^45^, promotes both the relocation of dysfunctional forks to NPCs and RDR efficiency in a Pli1-dependent manner^15^. This supports the notion that mono-SUMOylated or chain-free multi-SUMOylated factors are potential targets of a proteasome and Slx8-dependent pathway that ensures the progress of restarted DNA polymerases. SUMO chains-independent functions of STUBL are documented, including the relocation of forks collapsed at CAG repeats via mono-SUMOylation recognized by the SUMO interacting motif of ScSlx5^17^. The human STUBL RNF4 can also bind the substrate ETV4 mono-SUMOylated on multiple lysines, in a process requiring the multiple SIM domains of RFN4^46^.

Our work also identified that in the absence of the nuclear basket nucleoporin Alm1, the RFB was no longer enriched at the NP. To our knowledge, TPR homologs have not been involved in anchoring DNA lesions to NPCs in yeast models. Upon telomeric replication stress, human telomeres relocate to the NP and associate with NPC components, including TPR, to resolve replication defects^19^. Depletion of human TPR is associated with a variety of replication defects and TPR was proposed to coordinate at NPCs a network of factors involved in RNA metabolism to protect cells from RNA-mediated replication stress^47^. Given the nuclear morphology alterations in the absence of Alm1, we do not favor the hypothesis of a direct involvement of Alm1 in anchoring dysfunctional forks at NPCs. In human cells, the mobility of stressed forks towards the NP requires F-nuclear actin polymerization^8,20^, but such a mechanism has not been reported in yeasts. We estimated that, in the absence of Alm1, the RFB must explore a nuclear volume 40 % larger to reach the NP and associate with NPCs whose abundance is reduced by one quarter. We therefore favor the hypothesis that the lack of relocation is an indirect effect due to alterations of nuclear morphology and NPC density.

Overall, this work uncovers two mechanisms by which the NPC environment ensures the dynamic of HR-dependent replication restart, streamlining the need for dysfunctional forks to change nuclear positioning. Ulp1-associated NPCs contribute to the efficient initiation of restarted DNA synthesis to engage a Polδ/Polδ DNA synthesis, by counteracting the inhibitory effect of SUMO chains, whereas proteasome-associated NPCs foster the progression of restarted DNA synthesis, in a SUMO chains independent manner. These two functions cannot compensate for each other, are differently required and control distinct dynamics of replication resumption at dysfunctional forks. Moreover, our work indicates that multiple SUMOylated targets are likely engaged to limit HR-dependent DNA synthesis.

## Funding and Acknowledgements

The authors thank the Multimodal Imaging Center Imaging Facility of the Institut Curie - CNRS UMS2016 / Inserm US43 / Institut Curie / Université Paris- and the Flow Cytometry Facility of the Orsay site of Institut Curie. This study was supported by grants from the Institut Curie, the CNRS, the Fondation LIGUE contre le cancer “Equipe Labellisée 2020 (EL2020LNCC/Sal), the ANR grant NIRO (ANR-19-CE12-0023-01). KS has received a PhD fellowship from the Fondation LIGUE contre le cancer and a 4^th^-year PhD grant from Fondation ARC. KK was supported by the program “Excellence Initiative - Research University” for the University of Wrocław of the Ministry of Education and Science from Poland, under grant number IDN.CBNDR 0320/2020/20. AC acknowledges Wellcome grant 110047/Z/15/Z.

The funders had no role in study design, data collection and analysis, the decision to publish, or preparation of the manuscript.

## Author contributions

K.K., K.S., K.N., S.C. and K.F. performed the experiments.

K.K. K.S. A.C. and S.A.E.L contributed to experimental design and data analysis.

L.B. provided expertise to perform and analyze cell imaging.

K.N. and A.C provided the expertise to analyze Pu-seq data.

K.K, A.C., K.N., L.B. and S.A.E.L wrote the manuscript.

## Declaration of interests

The authors declare no competing interests.

## Methods

### Standard yeast genetics

Yeast strains used in this work are listed in Table S1. Gene deletion and tagging were performed by classical genetic techniques. To assess the sensitivity of chosen mutants to genotoxic agents, mid log-phase cells were serially diluted and spotted onto yeast extract agar plates containing hydroxyurea (HU), methyl methanesulfonate (MMS), campthotecin (CPT), bleomycin (bleo). Strains carrying the *RTS1* replication fork block sequence were grown in minimal medium EMMg (with glutamate as a nitrogen source) with addition of appropriate supplements and 60 µM thiamine (barrier inactive, OFF). The induction of replication fork block was obtained by washing away the thiamine and further incubation in a fresh medium for 24 hours (barrier active, ON).

### Live cell imaging

For snapshot microscopy, cells were grown in filtered EMMg with or without 60 µM thiamine for 24 hours to exponential phase (RFB OFF and RFB ON), then centrifuged and resuspended in 500 µL of fresh EMMg. 1 µL from the resulting solution was dropped onto Thermo Scientific slide (ER-201B-CE24) covered with a thin layer of 1.4 % agarose in filtered EMMg^15^. 21 z-stack pictures (each z step of 200 nm) were captured using a Nipkow Spinning Disk confocal system (Yokogawa CSU-X1-A1) mounted on a Nikon Eclipse Ti E inverted microscope, equipped with a 100x Apochromat TIRF oil-immersion objective (NA: 1.49) and captured on sCMOS Prime 95B camera (Photometrics) operated through MetaMorph^®^ software (Molecular Devices). GFP and m-Cherry proteins were excited with a 488 nm (Stradus® - Vortran Laser Technology, 150mW) and a 561 nm (Jive^TM^ - Cobolt, 100mW) lasers, respectively. A quad band dichroic mirror (405/488/568/647 nm, Semrock) was used in combination with single band-pass filters of 525/50 or 630/75 for the detection of GFP and m-Cherry, respectively. Fluorescence and bright-field 3D images were taken at every 0.3µm over a total of 4.5µm thickness. Exposure time for GFP channel was 500 ms, for mCherry 1000 ms. During the imaging, the microscope was set up at 25°C. For the experiment on Ulp1 and Cut11, the Gataca Live SR module (Müller et al., 2016, Gataca Systems), implemented on the Spinning Disk confocal system, was used to generate super-resolution images with lateral image resolution improvement (around 120 nm).

### Image analysis

Images were mounted and analyzed with Fiji software^48^. First, the 3D Z series are converted into 2D projection based on maximum intensity values. The quantification of Ulp1 and Cut11 was performed using a homemade macro. The user draws manually all nuclear rings on the merge images as a first step. Then automatically, 3 types of regions were created from the manual annotation:

- the nucleus was obtained by enlarging the manual annotation to? 3 pixels
- the nucleoplasm was obtained by shrinking the nucleus to? 8 pixels
- the nuclear periphery has been extracted from the previous two regions by selecting only those pixels that are not common.

Several measurements were exported for all regions, such as perimeter of nuclei in pixels, area in pixels², the fluorescence density of a protein (annotated as “Mean fluorescence intensity” in Fiji: this value represents the Raw Integrated Density measured in the selection and normalized by the area of the same selection) and the total fluorescence intensity of the protein (annotated as “RawIntDen”(Raw Integrated Density) in Fiji: this value represents the sum of all pixels intensities in the selection). To assess the co-localization of Ulp1 and Cut11 proteins, the JACoP plugin^49^ was used to study the correlation between the intensities of these two proteins in different NPC mutant strains. Pearson and Manders’ coefficients were calculated for each nucleus obtained previously. Before running the analysis, pre-processing was applied (background subtraction using the rolling ball algorithm with a radius of 20 pixels and a Gaussian filter (sigma 1)) to reduce image noise and facilitate detection of the Ulp1 and Cut11 proteins needed to calculate Manders’ coefficients. The “Default” thresholding method was used for the detection of Ulp1-GFP and Cut11-mCherry positive signals.

### 2DGE analysis of replication intermediates

Exponential cells (2.5×10^9^) were treated with 0.1% sodium azide and subsequently mixed with frozen EDTA (of final concentration at 80 mM). Genomic DNA was crosslinked with trimethyl psoralen (0.01 mg/mL, TMP, Sigma, T6137) added to cell suspensions for 5 min in the dark. Next, cells were irradiated with UV-A (365 nm) for 90 s at a constant flow 50 mM/cm^2^. Subsequently, cell lysis was performed by adding lysing enzymes (Sigma, L1412) at a concentration of 0.625 mg/mL and zymolyase 100 T (Amsbio, 120493-1) at 0.5 mg/mL. Obtained spheroplasts were next embedded into 1 % low melting agarose (InCert Agarose 50123, Lonza) plugs and incubated overnight at 55 °C in a digestion buffer with 1 mg/mL of proteinase K (Euromedex EU0090). Then plugs were washed with TE buffer (50 mM Tris, 10 mM EDTA) and stored at 4 °C. Digestion of DNA was performed using 60 units of restriction enzyme *Ase*I (NEB, R0526M) per plug. Samples were then treated with RNase (Roche, 11119915001) and beta-agarase (NEB, M0392L). Melted plugs were equilibrated to 0.3 M NaCl concentration. Replication intermediates were purified using BND cellulose (Sigma, B6385) poured into columns (Biorad, 731-1550)^50^. RIs were enriched in the presence of 1M NaCl 1.8% caffeine (Sigma, C-8960), precipitated with glycogen (Roche, 1090139001) and migrated in 0.35 % agarose gel (1xTBE) for the first dimension. The second dimension was cast in 0.9 % agarose gel (1xTBE) supplemented with EtBr. Next, DNA was transferred to a nylon membrane (Perkin-Elmer, NEF988001PK) in 10x SSC. Finally, membranes were incubated with ^32^P-radiolabeled *ura4* probe (TaKaRa *Bca*BEST^TM^ Labeling Kit, #6046 and alpha-^32^P dCTP, Perkin-Elmer, BLU013Z250UC) in Ultra-Hyb buffer (Invitrogen, AM8669) at 42°C. The signal of replication intermediates was collected in phosphor-imager software (Typhoon-trio) and quantified by densitometric analysis with ImageQuantTL software (GE healthcare). The ‘tail signal’ was normalized to the overall signal corresponding to arrested forks.

### Replication slippage assay

The frequency of *ura4+* revertants using the *ura4-sd20* allele was determined as follows. 5-FOA (EUROMEDEX, 1555) resistant colonies were grown on plates containing uracil with or without thiamine for 2 days at 30 °C and subsequently inoculated into EMMg supplemented with uracil for 24 h. Then cultures were diluted and plated on EMMg complete (for cell survival) and on EMMg without uracil, both supplemented with 60 µM thiamine. After 5-7 days of incubation at 30°C colonies were counted to determine the frequency of ura4+ reversion. To obtain the true occurrence of replication slippage by the *RTS1*-RFB, independently of the genetic background, we subtracted the replication slippage frequency of the strain devoid of RFB (considered as spontaneous frequency) from the frequency of the strain containing *the t-ura4sd20<ori* construct, upon expression of Rtf1.

### Flow cytometry

Flow cytometry analysis of DNA content was performed as follows^51^: cells were fixed in 70 % ethanol and washed with 50 mM sodium citrate, digested with RNAse A (Sigma, R5503) for 2 hours, stained with 1µM Sytox Green nucleic acid stain (Invitrogen, S7020) and subjected to flow cytometry using FACSCANTO II (BD Biosciences).

### Whole protein extract analysis

Aliquots of 1×10^8^ cells were collected and disrupted by bead beating in 1 mL of 20 % TCA (Sigma, T9159). Pellets of denatured proteins were washed with 1M Tris pH 8 and resuspended in 2x Laemmli buffer (62.5 mM Tris pH 6.8, 20 % glycerol, 2 % SDS, 5 % β-mercaptoethanol with bromophenol blue). Samples were boiled before being subjected to SDS-PAGE on Mini-PROTEAN TGX Precast Gel 4-15 % (Biorad, 4561086). Western blot using anti-GFP (Roche, 11814460001), anti-HA (Santa Cruz Biotechnology, sc-57592), anti-TIR1 (MBL, PD048), anti-PCNA (Santa Cruz, sc-56) or anti-tubulin (Abcam, Ab6160) antibodies was performed. For the analysis of cellular patterns of global SUMOylation, whole protein extraction was performed as follows: aliquots of 2×10^8^ cells were collected and resuspended in 400µl of water. The cell suspensions were mixed with 350 µl of freshly prepared lysis buffer (2M NaOH, 7% β-mercaptoethanol) and 350µl of 50% TCA (Sigma, T9159). After spin, pellets were further washed with 1M Tris pH 8 and resuspended in 2x Laemmli buffer (62.5 mM Tris pH 6.8, 20 % glycerol, 2 % SDS, 5 % β-mercaptoethanol with bromophenol blue). Samples were boiled before being subjected to SDS-PAGE on Mini-PROTEAN TGX Precast Gel 4-15 % (Biorad, 4561086). Western blot using anti-SUMO antibody (non-commercial, produced in rabbit by Agro-Bio) was performed.

### Pulse field gel electrophoresis

Yeast cultures were grown to logarithmic phase in rich YES medium to a concentration of 5 x 10^6^/mL, synchronized in 20 mM HU for 4 hours, and subsequently released to fresh YES medium. At each time point, 20 mL of cell culture was harvested, washed with cold 50 mM EDTA pH 8 and digested with lyticase (Sigma, L4025) in CSE buffer (20 mM citrate/phosphate pH 5.6, 1.2 M sorbitol, 40 mM EDTA pH 8). Next cells were embedded into 1% UltraPure^TM^ Agarose (Invitrogen, 16500) and distributed into 5 identical agarose plugs for each time point. Plugs were then digested with Lysis Buffer 1, LB1 (50 mM Tris-HCl pH 7.5, 250 mM EDTA pH 8, 1 % SDS) for 1.5 hours at 55°C and then transferred to Lysis Buffer 2, LB2 (1 % N-lauryl sarcosine, 0.5 M EDTA pH 9.5, 0.5 mg/mL proteinase K) o/n at 55°C. The next day, LB2 was changed to a fresh one and digestion was continued o/n at 55°C. After, plugs were kept at 4°C. To visualize intact chromosomes, one set of plugs was run on a Biorad CHEF-DR-III pulse field gel electrophoresis (PFGE) system for 60 h at 2.0 V/cm, angle 120°, 14°C, 1800 s single switch time, pump speed 70 in 1x TAE buffer. Separated chromosomes were stained in ethidium bromide (10 μg/mL) for 30 min, washed briefly in 1x TAE and visualized with a UV trans-illuminator.

### Pu-Seq

The published protocol^52^ was used with minor modifications: size selection was performed using a Blue Pippin (Sage Science). We used *rnh201-RED* instead of *rnh201::kan*^26^. Sequence files were aligned with Bowtie2 and alignment data converted to counts with custom Perl script^52^. Analysis of polymerase usage was performed with custom R script^52^. Sequence data is available under GEO dataset GSE247371.

## STATISTICAL ANALYSIS

Quantitative densitometric analysis of Southern blots after 2DGE was carried out using ImageQuant software. The ‘tail signal’ of resected forks was normalized to the overall signal corresponding to arrested forks.

Quantification of PFGE was performed using ImageJ and presented as % of migrating chromosomes relative to asynchronous profile. Cell imaging was performed using METAMORPH software and processed and analyzed using ImageJ software^48^. The explanation and definitions of values and error bars are mentioned within the figure legends. In most experiments, the number of samples is > 3 and obtained from independent experiments to ensure biological reproducibility. For all experiments based on the analysis of cell imaging, the number of nuclei analyzed is mentioned in the figure legends. Statistical analysis was carried out using Mann-Whitney U tests, Student’s *t*-test and Fischer’s exact test.

## DATA AVAILABILITY

The source data files have been deposited to Mendeley data and are available at “Schirmeisen, Naiman et al 2023”, Mendeley Data, V1, doi: 10.17632/2kgnb9d66r.1. The source data underlying Figs 2a, 2c-d, 2b-d, 2g, 3a-b, 4b-c, 4e, 5a-c, and Supplementary Figs 1a, 1c-d, 2a, 2c, 3a-d, 4a-d are provided as a Source Data file. RAW data from Pu-seq experiments are available under GEO dataset GSE247371. All relevant data are available and further information and requests for reagents and resources should be directed to and will be fulfilled by Dr. Sarah A.E. Lambert.

## Supplemental Figure

**Figure S1:**
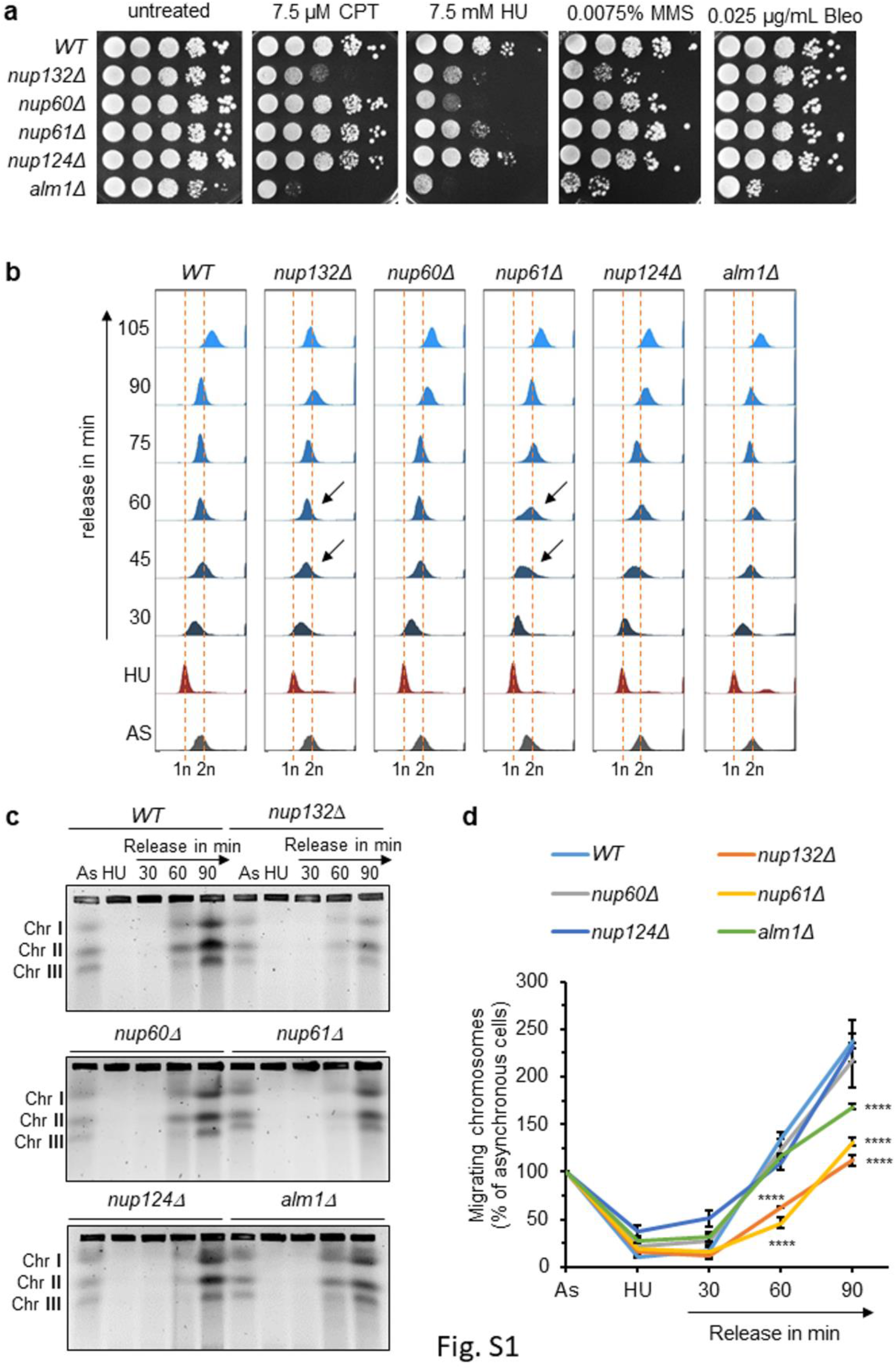
Role of the nuclear basket in the recovery from HU-induced stalled forks. **a**. Sensitivity of indicated strains to indicated genotoxic drugs. Ten-fold serial dilutions of exponential cultures were dropped on appropriate plates. Bleo: bleomycin; CPT: camptothecin; HU: hydroxyurea; MMS: methyl methane sulfonate. **b.** Flow cytometry analysis of indicated strains in indicated conditions. Logarithmically growing cells (AS: Asynchronous cells) were exposed to 20 mM HU for 4 hours (HU time point) and then released into fresh, HU-free, rich medium YES at 30°C to monitor S-phase progression at the indicated time after release. Arrows indicate the delay in S phase progression compared to *WT*. **c**. Analysis of chromosomes by pulse field gel electrophoresis (PFGE) in the above-mentioned conditions (as in b). Representative images of chromosome migration during PFGE in indicated strains and conditions. **d.** Quantification of % of chromosomes migrating into the gel after release from HU block. Values are means of two independent biological replicates ± standard deviation (SD). *p* value was calculated by two-sided Fisher’s exact test (**** *p* ≤0.0001).

**Figure S2:**
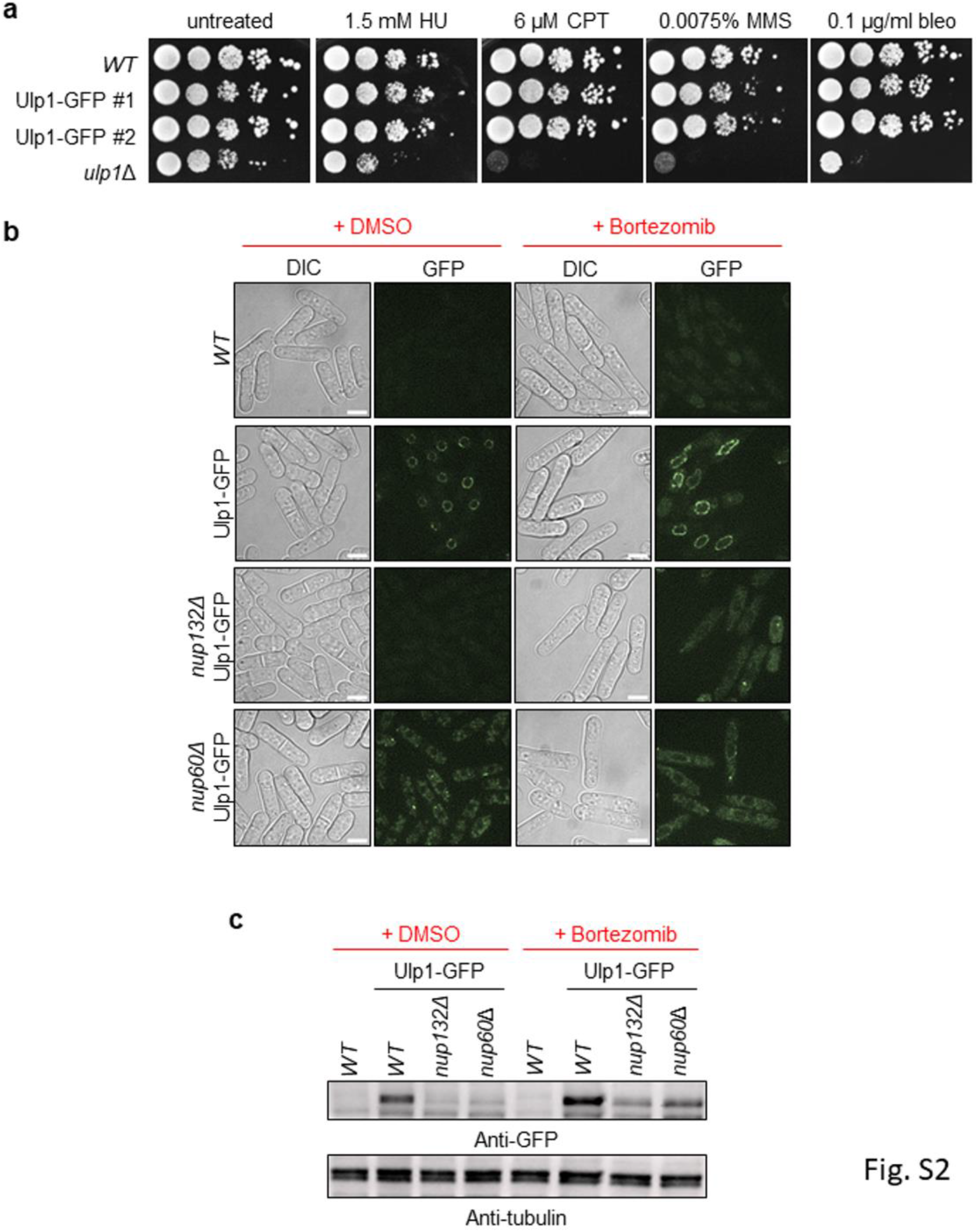
The downregulation of Ulp1 expression is caused by the proteasome. **a**. Ulp1-GFP is a functional fusion protein. Ten-fold serial dilutions of exponential cultures were dropped on appropriate plates. Bleo: bleomycin; CPT: camptothecin; HU: hydroxyurea; MMS: methyl methane sulfonate. **b**. Cell imaging of Ulp1-GFP in indicated strains and conditions. Representative cell images of Ulp1-GFP in indicated strains in presence or absence of bortezomib. Scale bar: 5µm. **c.** Expression of Ulp1-GFP in indicated strains and conditions. An untagged WT strain was included as a control for antibody specificity. Tubulin was used as a loading control.

**Figure S3:**
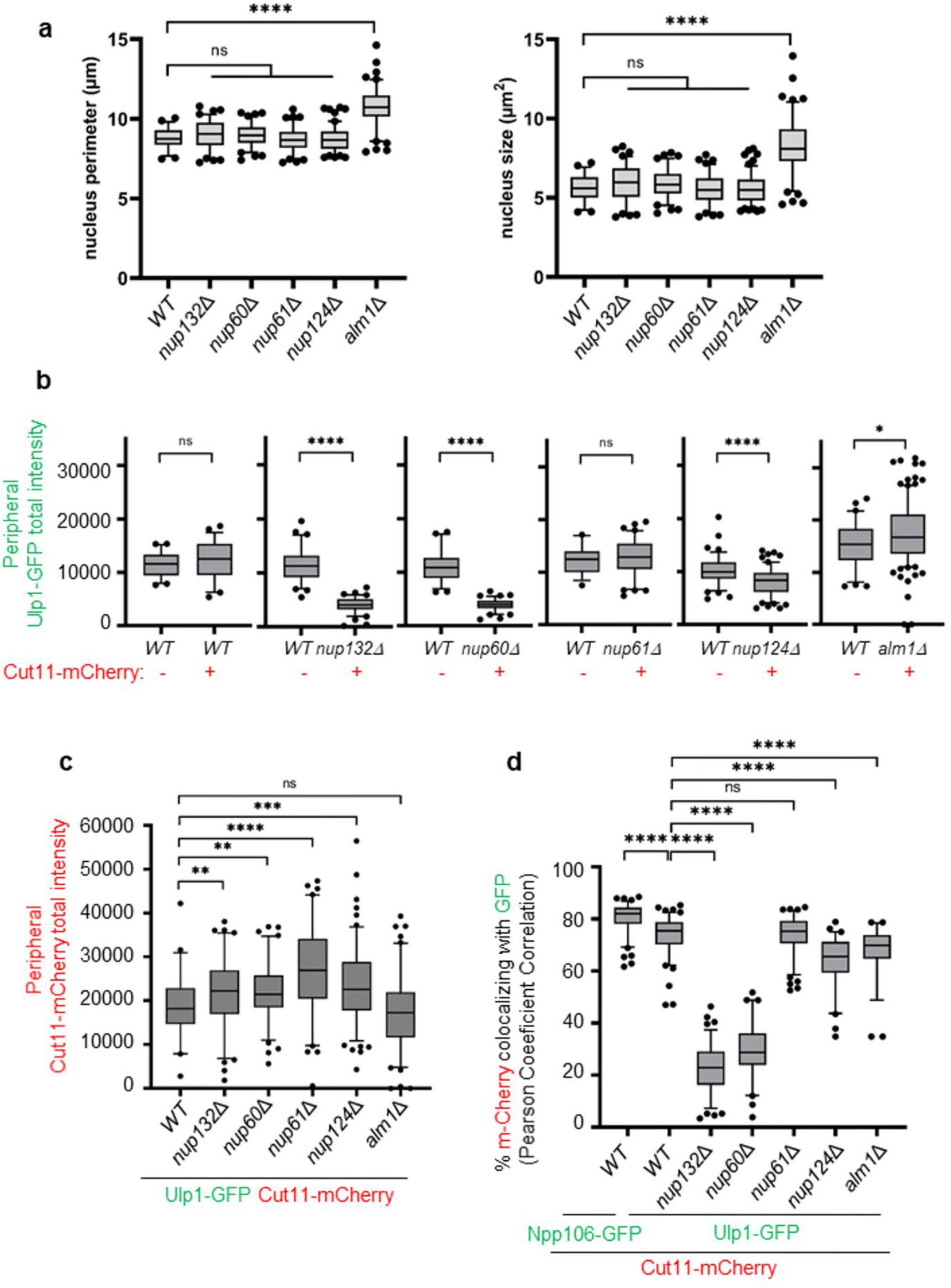
Image quantification of nuclear morphology parameters, Ulp1-GFP and Cut11-mCherry intensity. **a.** Box-and-whisker plots of nucleus perimeter (left panel) and nucleus size (right panel) in indicated strains. Boxes represent the 25/75 percentile, black lines indicate the median, the whiskers indicate the 5/95 percentile and dots correspond to minimum and maximum values. *p* value was calculated by Mann-Whitney U test (**** *p* ≤0.0001; ns: non-significant). At least 50 nuclei were analyzed for each strain. **b.** Box-and-whisker plots of Ulp1-GFP total intensity (raw integrated density) at the nuclear periphery in indicated strains and conditions. Boxes represent the 25/75 percentile, black lines indicate the median, the whiskers indicate the 5/95 percentile and dots correspond to minimum and maximum values. *p* value was calculated by Mann-Whitney U test (**** *p* ≤0.0001; * p≤0.05; ns: non-significant). At least 50 nuclei were analyzed for each strain. **c.** Box-and-whisker plots of Cut11-mCherry total intensity (raw integrated density) at the nuclear periphery in indicated strains and conditions. Boxes represent the 25/75 percentile, black lines indicate the median, the whiskers indicate the 5/95 percentile and dots correspond to minimum and maximum values. *p* value was calculated by Mann-Whitney U test (**** *p* ≤0.0001; *** p≤0.001; ** p≤0.01; ns: non-significant). At least 50 nuclei were analyzed for each strain. **d.** Box-and-whisker plots of co-localization between Cut11-mCherry and Ulp1-GFP (using the Pearson correlation coefficient) in indicated strains. The co-localization between the Npp106-GFP, an inner ring nucleoporin of NPC, and Cut11-mCherry, was performed as a control to show maximum correlation between intensities of both proteins at the resolution achieved on the images. Boxes represent the 25/75 percentile, black lines indicate the median, the whiskers indicate the 5/95 percentile and dots correspond to minimum and maximum values. *p* value was calculated by Mann-Whitney U test (**** *p* ≤0.0001; ns: non-significant). At least 50 nuclei were analyzed for each strain.

**Figure S4:**
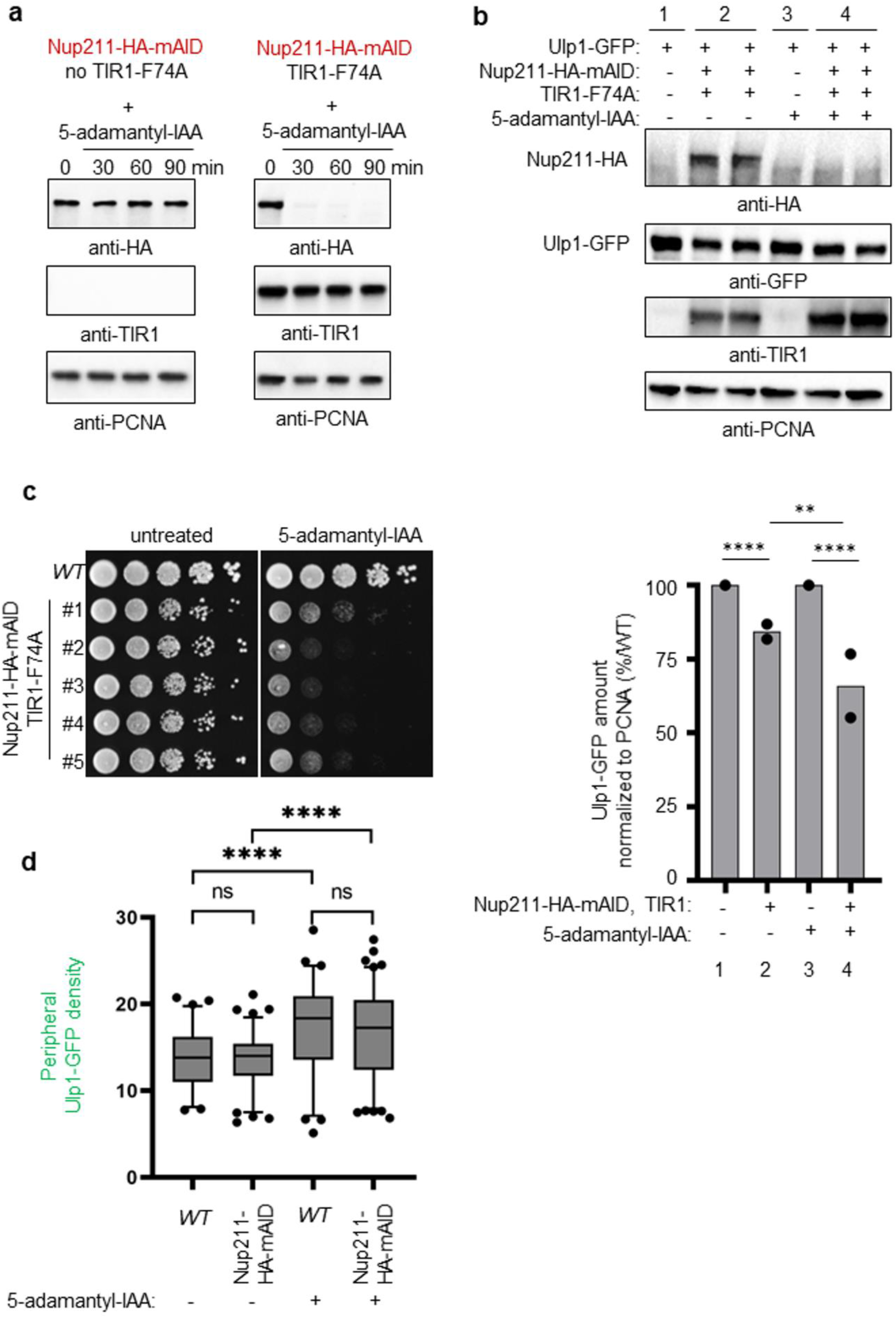
An auxin-induced degron approach to conditionally downregulate Nup211. **a.** Expression of Nup211-HA-mAID fusion protein in strains expressing TIR1-F74A (right panels) or not (left panels) as a function of time (in minute) upon addition of 5-adamentyl-IAA. PCNA was used as a loading control. **b.** Top panels: expression of Ulp1-GFP, Nup211-HA-mAID and TIR1in indicated conditions. PCNA was used as loading control. Bottom panel: quantification. Dots represent values obtained from independent biological experiments. The normalized amount of Ulp1 was calculated by dividing the GFP signal by PCNA. The normalized amount of Ulp1-GFP in mutants was indicated as a percentage of the *WT*. *p* value was calculated by two-sided Fisher’s exact test (**** p≤0.0001; ** p≤0.01). **c**. Cell growth assay of indicated strains. Ten-fold serial dilutions of exponential cultures were dropped on plates containing 5-adamantyl-IAA (right panel) or not (left panel). Five independent clones expressing Nup211-HA-mAID were tested alongside the *WT* strain. Note the cell growth defect of Nup211-HA-mAID strains in the absence of 5-adamantyl-IAA is indicative of a lack of Nup211 functionality. **d.** Box-and-whisker plots of Ulp1-GFP density (mean fluorescence intensity) at the nuclear periphery in indicated strains and conditions. Boxes represent the 25/75 percentile, black lines indicate the median, the whiskers indicate the 5/95 percentile and dots correspond to minimum and maximum values. *p* value was calculated by Mann-Whitney U test (**** *p* ≤0.0001; ns: non-significant). At least 50 nuclei were analyzed for each strain.

**Supplementary Table 1.**
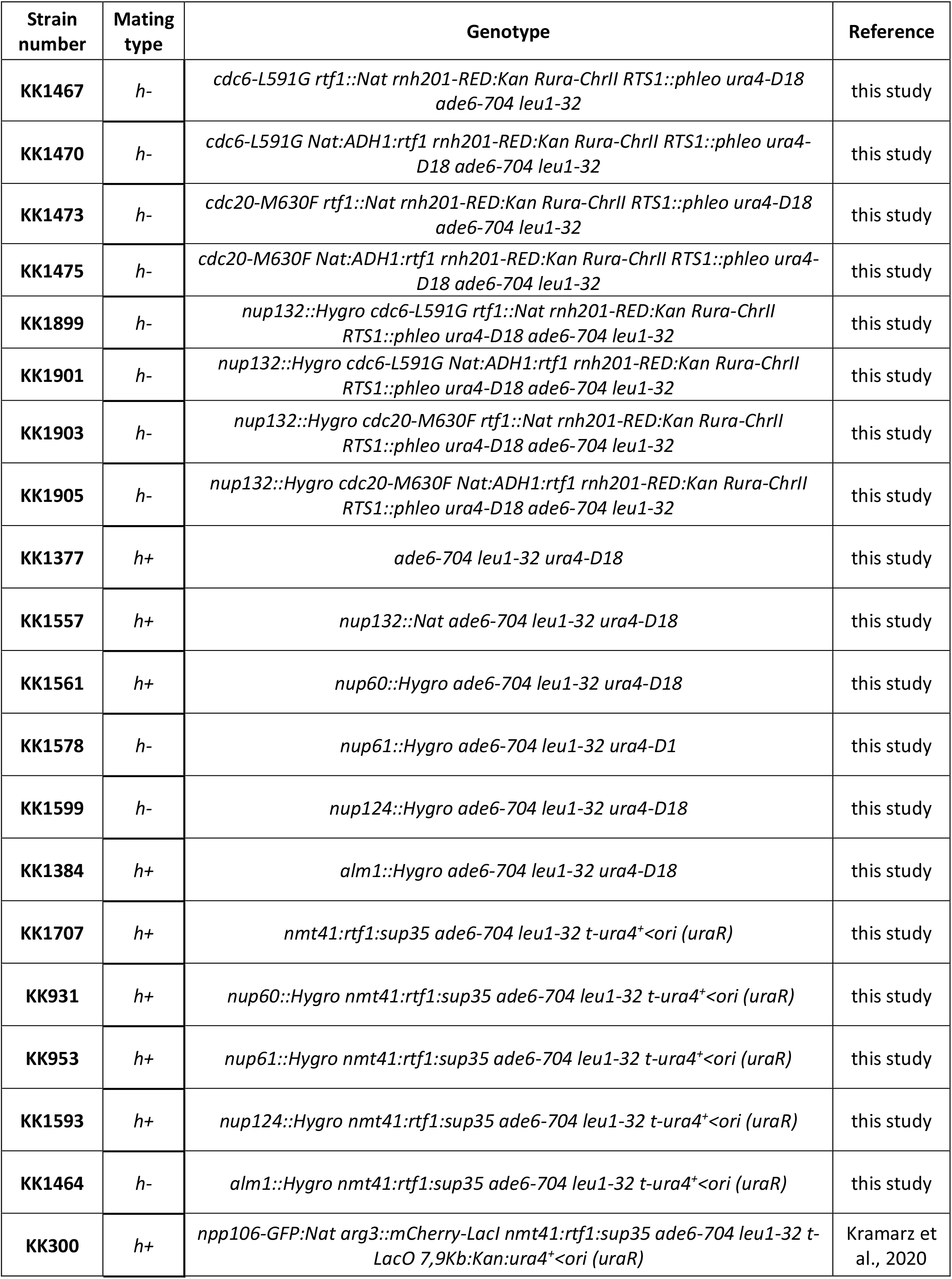
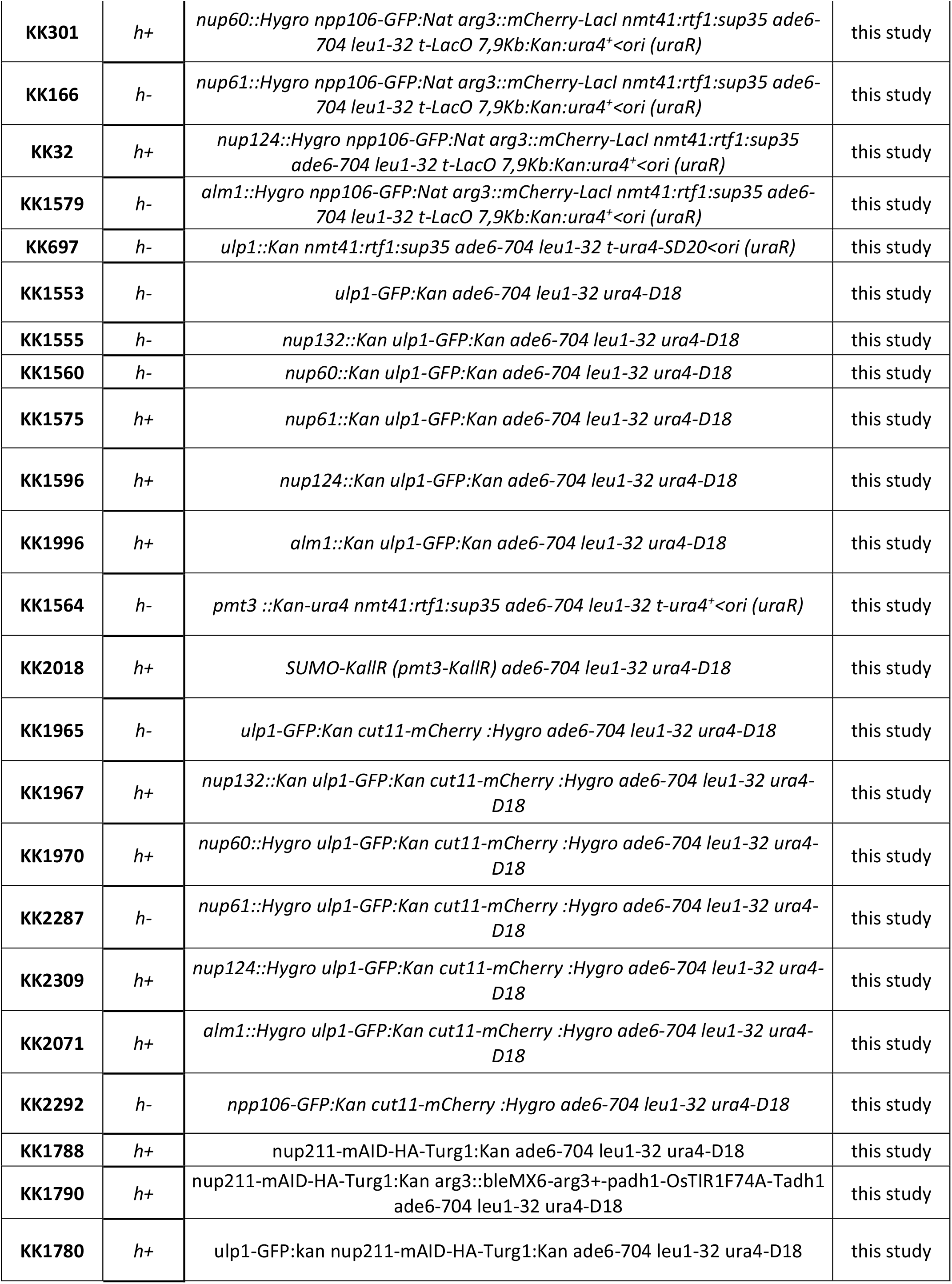
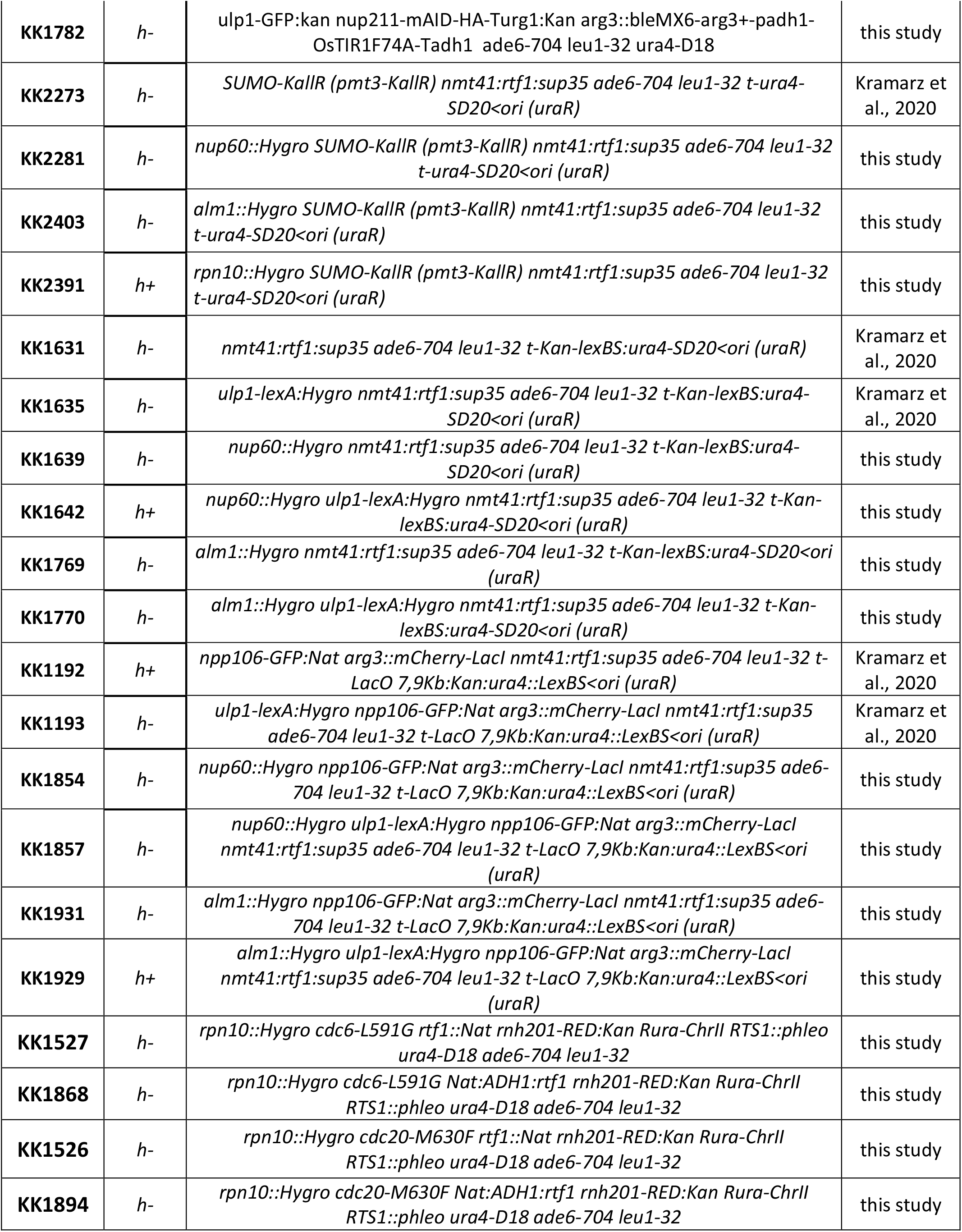

## Notes

### Competing Interest Statement

The authors have declared no competing interest.

